# Resistance to abemaciclib is associated with increased metastatic potential and lysosomal protein deregulation in breast cancer cells

**DOI:** 10.1101/2023.04.17.537215

**Authors:** Erin R. Scheidemann, Diane M. Demas, Chunyan Hou, Junfeng Ma, Wei He, Katherine N. Weilbaecher, Ayesha N. Shajahan-Haq

## Abstract

Cyclin dependent kinase 4 and 6 inhibitors (CDK4/6i) such as abemaciclib are routinely used to treat metastatic estrogen receptor positive (ER+)/HER2-negative breast cancer. However, adaptive mechanisms inhibit their effectiveness and allow for disease progression. Using murine metastatic ER+ breast cancer cells, we show that acquired resistance to abemaciclib is accompanied by increase in metastatic potential. Mass spectrometry-based proteomics from abemaciclib sensitive and resistant cells showed that lysosomal proteins including CTSD (cathepsin D), CTSA (cathepsin A) and CD68 were significantly increased in resistant cells. Combination of abemaciclib and a lysosomal destabilizer, such as hydroxychloroquine (HCQ) or bafilomycin A1, re-sensitized resistant cells to abemaciclib. Also, combination of abemaciclib and HCQ decreased migration and invasive potential and increased lysosomal membrane permeability (LMP) in resistant cells. Pro-survival BCL2 protein levels were elevated in resistant cells, and a triple treatment with abemaciclib, HCQ, and BCL2 inhibitor, venetoclax, significantly inhibited cell growth compared to treatment with abemaciclib and HCQ. Furthermore, resistant cells showed increased levels of TFEB (Transcription Factor EB), a master regulator of lysosomal-autophagy genes, and siRNA mediated knockdown of *TFEB* decreased invasion in resistant cells. *TFEB* gene was found to be mutated in a subset of invasive human breast cancer samples, and overall survival analysis in ER+, lymph node-positive breast cancer showed that increased *TFEB* expression correlated with decreased survival. Collectively, we show that prolonged exposure to abemaciclib in ER+ breast cancer cells leads to resistance accompanied by an aggressive phenotype that is partly supported by deregulated lysosomal function.

**Implications**: Our data implicate that resistance to abemaciclib is associated with deregulation of lysosomes and augmented metastatic potential, and therefore, the lysosomal pathway could be a therapeutic target in advanced ER+ breast cancer.

**GRAPHICAL ABSTRACT:** 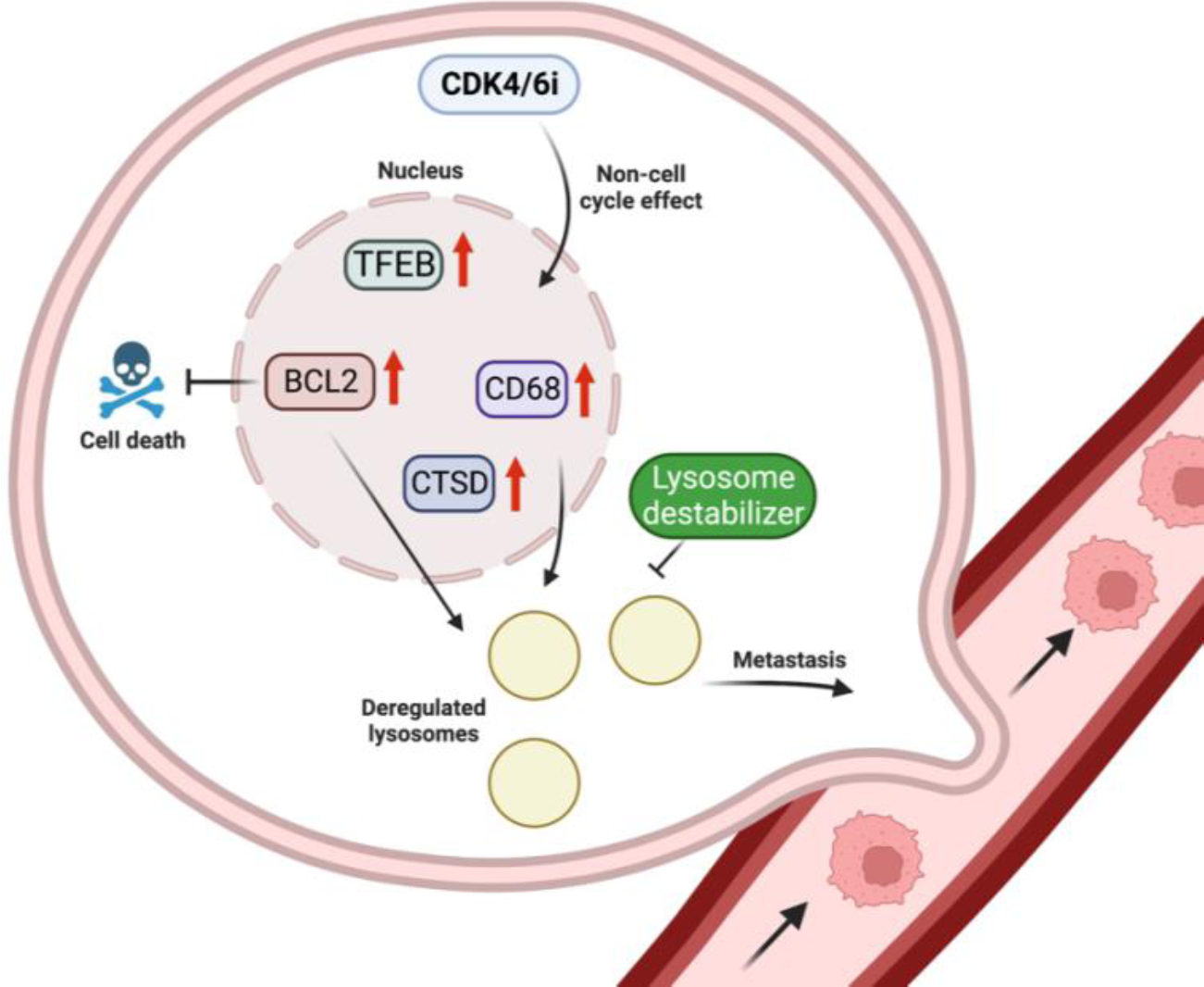

---

<img/>

## INTRODUCTION

Resistance to CDK4/6 inhibitors (CDK4/6i) is a current clinical problem in treating hormone receptor positive (HR+)/HER2-breast cancer. Three CDK4/6i are currently FDA approved: palbociclib, ribociclib, and abemaciclib. These drugs inhibit cell cycle dependent kinases, CDK4 and CDK6, from binding with Cyclin D1 to prevent RB1 phosphorylation and induce G1 arrest. It is estimated that 20% of patients with tumors resistant to endocrine therapy have intrinsic resistance to CDK4/6i, and all patients eventually go on to acquire resistance to these drugs over time (1,2). Currently, there is no drug specifically intended for use in CDK4/6i resistant breast cancer. Abemaciclib is the only CDK4/6i currently approved as a monotherapy in metastatic breast cancer (3), and has been recently been approved for early high-risk breast cancer in combination with endocrine therapy (4). Abemaciclib is 5-times more potent against CDK4 than palbociclib or ribociclib (5) and can cross the blood brain barrier (6). Its relatively weak activity against CDK6 adds to its clinical value, as patients do not experience the CDK6 specific-side effect of neutropenia, and thus, no drug holiday is needed.

In the nucleus, CDK4/6 phosphorylates Transcription Factor EB (TFEB) and Transcription Factor Binding to IGHM Enhancer 3 (TFE3), master regulators of lysosomal-autophagy pathway genes. Phosphorylation of TFEB/TFE3 causes them to be shuttled to the cytoplasm, and thus TFEB/TFE3-mediated gene transcription of the Coordinated Lysosomal Expression and Regulation (CLEAR) gene network is prevented (7). TFEB/TFE3 and the CLEAR network tightly regulate the activity of many enzymes to control lysosomal functions such as exocytosis and autophagy. Abemaciclib mediated inhibition of CDK4/6 can prevent nuclear phosphorylation of TFEB/TFE3 and increase lysosomal biomass and function, which mediates a novel lysosomal-mediated form of cell death in sensitive cells (7,8). In a subset of triple negative breast cancer (TNBC) cells with intact RB1, TFEB mediated transcription of lysosomal genes was shown to increase in lysosomal mass and enabled these TNBC cells to intrinsically resist palbociclib by sequestering the drugs into the lysosomes (9). However, the mechanism by which ER+ breast cancer cells use lysosomes to acquire resistance to long-term CDK4/6i treatment remains unknown.

The relationship between CDK4/6 inhibition and lysosomal mass/function is particularly interesting in the context of advanced disease. Though drug resistance is often seen in the advanced setting, only recently has research started to investigate connections between drug resistance and cancer metastasis (10,11). Lysosomes have been implicated in metastasis in multiple ways. The surface of highly metastatic cancer cells has increased expression of lysosomal-associated membrane protein 1 (LAMP1) (12). Likewise, cathepsins, proteolytic lysosomal enzymes, can degrade the extracellular matrix (ECM) to facilitate invasion and thus have been associated with metastasis (13–16). Given this evidence and the recent line of thought that drug resistance is directly linked to metastasis, we hypothesized that deregulated activity in abemaciclib resistant cells causes cells to gain more migratory/invasive abilities. In this study, we used ER+ murine and human breast cancer cells with acquired resistance to abemaciclib to show that resistant cells with altered lysosomal proteins have increased metastatic behavior *in vitro*. As a result, combination of a lysosome-targeting drug and abemaciclib sensitizes resistant cells by inducing lysosome membrane permeabilization (LMP).

## MATERIALS AND METHODS

### Reagents

Abemaciclib (LY2835219), hydroxychloroquine (NSC 4375), and bafilomycin A1 (NSC 381866) were purchased from Cayman Chemical Company (Ann Arbor, MI). Palbociclib (PD0332991) and ribociclib (LEE011) were purchased from Selleck Chemicals (Houston, TX). All other reagents were purchased from Sigma Aldrich (St Louis, MO). Abemaciclib was dissolved in ethanol (EtOH). All other drugs were dissolved in dimethyl sulfoxide (DMSO). For in vitro assays, negative control was 0.02% ethanol or DMSO.

### Cell culture

Murine C57BL/6 PyMT-B6 and PyMT-Bo1 ER+ breast cancer cell lines were established as described previously (17). PyMT-Bo1 cells were derived from bone metastasis of PyMT-B6 tumors *in vivo*. MCF7 and LCC1 cells were obtained from Georgetown University Medical Center Tissue Culture Shared Resource. LCC1 cells were originally established from MCF7 cells and are estrogen-independent (18). Abemaciclib, palbociclib, and ribociclib-resistant cell lines were generated from PyMT-B6, PyMT-Bo1, and LCC1 cells in respective complete media supplemented with 1μM abemaciclib, palbociclib, or ribociclib for 6+ months. PyMT cells were cultured in Dulbecco’s Modified Eagle Medium (DMEM; Thermo Fisher Scientific, Waltham, MA) and supplemented with 10% fetal bovine serum (FBS; Life Technologies, Grand Island, NY) and penicillin-streptomycin (Thermo Fisher Scientific, Waltham, MA). MCF7 cells were cultured in Improved Minimum Essential Medium (IMEM; Thermo Fisher Scientific, Waltham, MA) supplemented with 10% charcoal-stripped calf serum (CCS; Life Technologies, Grand Island, NY) and 10% estradiol (19). LCC1 cells were cultured in Improved Minimum Essential Medium (IMEM; Thermo Fisher Scientific, Waltham, MA) supplemented with 5% charcoal-stripped calf serum (CCS; Life Technologies, Grand Island, NY). All cell lines were maintained in a humidified atmosphere with 5% CO_2_ at 37°C. All cells were authenticated by DNA fingerprinting and regularly tested for Mycoplasma infection.

### Cell proliferation assay

Murine PyMT cells were seeded with 1,000 cells per well in a 96-well plastic tissue culture plate per cell line. The first day after plating, cells were dosed with 0.02% vehicle (ethanol or DMSO) or various concentrations of the indicated drugs. These plates ended at 72 h. LCC1 cells were seeded with 3,500-10,000 cells per well and ended at 6 days. For crystal violet staining, plates were rinsed once with 1×PBS to remove cellular debris. Then, 100μL crystal violet was added to each well and rocked at room temperature for 1 h. To remove the stain, each plate was rinsed 5-10× with H_2_O. The plates were left at room temperature to dry overnight, then were rehydrated with 100μL 0.1M sodium citrate buffer in 50% ethanol. Plates were measured using a Vmax kinetic microplate reader (Molecular Devices Corp., Menlo Park, CA) with an absorbance of 560nM. Results were normalized to vehicle.

### Western blotting

Cells were washed once with cold 1×PBS. They were then lysed in radio-immunoprecipitation assay (RIPA) buffer supplemented with PhosSTOP phosphatase and CompleteMini protease inhibitors (Roche, Switzerland) for protein extraction. Proteins were separated by polyacrylamide gel electrophoresis (PAGE) using 4-12% gradient gels followed by protein transfer onto nitrocellulose membranes with iBLOT2 (Thermo Fisher Scientific, Waltham, MA). Membranes were blocked in 5% nonfat dry milk made in Tris-buffered saline with Tween-20 (TBST) and rocked at 4°C with the respective primary antibodies overnight. Secondary antibodies conjugated with horseradish peroxidase and SuperSignal West Dura Extended Duration Substrate (Thermo Fisher Scientific, Waltham, MA) were used to detect proteins of interest. The following antibodies were purchased from Cell Signaling Technology (Danvers, MA): CDK4 (#12790), CDK6 (#3136), phospho-RB1 (S780) (#3590), and Cyclin D1 (#2978), LC3B (#2775), Cleaved PARP (#9548S), BCL2 (#3498T), TFEB (#83010S), and CTSD for human cell lysates (#2284). ER alpha (#ab108398) and RB1 (#ab181616) antibodies were purchased from Abcam. P62 (#610832) was purchased from BD Biosciences. **β**-tubulin (#T7816) was acquired from Sigma-Aldrich. Actin (#sc-47778) antibody was purchased from Santa Cruz Biotech (Dallas, TX). CSTA for human cell lysates (#15020-1-AP) was purchased from Proteintech. CTSD for mouse cell lysates (#MAB1123-SP) and CTSA for mouse cell lysates (AF1029-SP) were bought from R&D Systems.

### Migration and Invasion Assays

For migration assays, 1×10^4^ cells in 100μL media were plated in the top of the Corning filter membrane insert (VWR, Radnor, PA) sitting inside of a 24 well tissue culture plate. For invasion assays, 0.5mL warm serum free media (SFM) was added on top of the Corning GFR Matrigel inserts (VWR, Radnor, PA). Matrigel was allowed to rehydrate for 2 h at 37°C. Then, the media was carefully removed so as to not disturb the matrigel. 1×10^4^ cells in 100μL media were plated in the top of the matrigel insert. Both assays used the same protocol after this point. Plates were incubated at 37°C for 10 minutes. Next, complete media was added into the lower chamber of the well. For invasion assays with TFEB siRNA, the protocol remained the same except both the cells in SFM and the complete media at the bottom of the well contained 20nM TFEB siRNA. Plates were incubated for 24 h at 37°C. Inserts were removed, and excess cells that did not move through the membrane/matrigel were gently removed from the surface using a cotton-tipped applicator. A Differential Quik III Stain Kit (Electron Microscopy Sciences, Hatfield, PA) was used to stain the cells trapped in the membrane or matrigel. Migration or invasion were quantitated by counting 10 fields at a magnification of 10× using a brightfield microscope. Each experiment was repeated in triplicate.

### Mass spectrometry-based proteomic analysis

PyMT-B6, PyMT-B6-abemaR, PyMT-Bo1 and PyMT-Bo1-abemaR cells were plated and grown in 10cm^2^ dishes in complete growth medium to 70% confluence. MCF7 cells were plated in 10 cm^2^ dishes for 24 h and treated with vehicle (0.02% EtOH) alone or 1 μM abemaciclib for 72 h. Four biological replicates were used for all cell lines. For proteomics analysis, cells were washed 3× with cold PBS, scraped down, centrifuged at 4°C followed by cold PBS wash for another 3 times. Equal amount of proteins from each sample were processed for proteomics, with a procedure published previously (20). In brief, samples resuspended in 5%SDS buffer were reduced with DTT and alkylated, followed by digestion on a S-Trap column (ProtiFi, LLC)with sequencing-grade Lys-C/trypsin (Promega). The resulting peptides were were analyzed with a nanoAcquity UPLC system (Waters) coupled with Orbitrap Fusion Lumos mass spectrometer (Thermo Fisher), with a procedure published previously (20). All data were acquired with data independent acquisition (DIA) mode. DIA data files were processed as described previously (20). In brief, the Spectronaut software version 15 (Biognosys) and a hybrid library were used (with default settings). Proteins identified and quantified were exported for comparison. Unpaired two-tailed Student’s t-test was used to define significant differentially expressed proteins (with fold changes >1.5 or <0.67; Q values < 0.05).

### Growth of primary tumors *in vivo*

PyMT-B6, PyMT-B6-abemaR, PyMT-Bo1 or PyMT-Bo1-abemaR cells (1×10^5^) were suspended in Matrigel (BD Biosciences) and orthotopically injected into the into the mammary fat pad of 8-week-old female C57BL/6 mice (n=5) (Charles River). Body weight and tumor size was measured twice a week for 2 weeks, at which point, primary tumors were excised and processed for histology. Hematoxylin and eosin (H&E) stained slides were evaluated by a veterinary pathologist at our Histopathology and Tissue Shared Resource.

### Transfection with *TFEB* siRNA

Cells were plated in 6 well (for protein assessment) or 96-well (for proliferation assay) plates. TFEB siRNAs (20nM, a mixture of 4 siRNA) or scrambled negative control were purchased from Dharmacon, Inc. (Lafayette, CO). siRNAs were transfected into the cells using RNA iMAX (ThermoFisher Scientific, Waltham, MA). Any indicated treatments were added 24 h post-transfection. Cells were lysed or plates were ended 72 h post-transfection. siRNA targeted the following sequences of TFEB: ‘CUACGAAGAUGAUGAAUAC’ ‘CAAGUUUGCUGCCCACGUG’ ‘GCAGAUGCCUAACACGCUG’ ‘ACUUAAUGCUCCUAGAUGA’.

### Cell Cycle and Cell Death Assays

<u>Cell cycle</u>: Cells were plated at 1 × 10^5^/well in 10cm^2^ dishes. Cells for time=0 h were grown in complete media and collected the following day. Treatment cells were grown in complete growth medium for 24 h before being treated with the indicated drug. Cells were then fixed in ethanol and analyzed by the Flow Cytometry Shared Resource according to the method of Vindelov et al. (21). <u>Apoptosis</u>: For the apoptosis assay, 1 × 10^5^/well cells were plated in 10cm^2^ dishes and treated with vehicle, 1μM abemaciclib, 5μM HCQ, or the combination for 72 h. Cells were stained with Annexin V-fluorescein isothiocyanate and propidium iodide, respectively (Thermofisher Scientific, Waltham, MA) according to the manufacturer’s protocol and fluorescence was measured by the Flow Cytometry Shared Resource at Georgetown University Medical Center. Each experiment was repeated at least three times. <u>Autophagy</u>: For the autophagy assay, 5×10^4^ cells were plated in 12 well plates and treated with vehicle, 1μM abemaciclib, 5μM HCQ, or the combination for 72 h. Starvation control cells were grown in SFM for 72 h. Cells were stained with CYTO-ID green detection reagent (Enzo Life Sciences, Farmingdale, NY) according to the manufacturer’s protocol and fluorescence was measured by the Flow Cytometry Shared Resource. Each experiment was repeated at least three times.

### LysoTracker studies for detecting lysosomal membrane permeability (LMP)

<u>Confocal Microscopy</u>: 12×10^4^ cells were plated in Ibidi µ-Slide 8 well glass bottom coverslips and treated with vehicle, 1μM abemaciclib, 5μM HCQ, or the combination for 24 h. Live cells were treated with LysoTracker Green DND-26 (Cell Signaling Technology, Danvers, MA) and Hoechst 33342 (Cell Signaling Technology, Danvers, MA) according to the manufacturer’s protocol and imaged with a Leica SP8 Confocal microscope through the Microscopy and Imaging Shared Resource.

### CD68 staining

Cells were washed twice with PBS and resuspended in Fixation and Permeabilization solution (BD Biosciences #554722) and incubated at 4°C for 20 minutes. Cells were washed twice with 1× Perm/Wash buffer (BD Biosciences #554723) and resuspended in 1× Perm/Wash buffer. TruStain FcX (BioLegend #101319) was added and cells were incubated on ice for 15 minutes. CD68 antibody (Biolegend #137013) was added and cells were incubated on ice for 30 minutes. Cells were washed with 1× Perm/Wash buffer and resuspended in Maxpar Cell Staining buffer (Fluidigm Corp #201068) for analysis by the Flow Cytometry Shared Resource.

### Analysis of *TFEB* gene mutation with cBioPortal

Presence of TFEB gene mutations in breast cancer was determined by using the cBio Cancer Genomics Potal (http://cbioportal.org) (22). A mutational analysis was performed in 7,298 breast cancer samples from nine studies: The Metastatic Breast Cancer Project (Provisional, December 2021), Proteogenomic landscape of breast cancer (CPTAC, Cell 2020), The Metastatic Breast Cancer Project (Archived, 2020), Metastatic Breast Cancer (INSERM, PLoS Med 2016), Breast Invasive Carcinoma (TCGA, Firehose Legacy), Breast Invasive Carcinoma (TCGA, Cell 2015), Breast Invasive Carcinoma (TCGA, PanCancer Atlas), Breast Cancer (METABRIC, Nature 2012 & Nat Commun 2016) and Breast Invasive Carcinoma (TCGA, Nature 2012).

### Statistical analysis, determination of synergistic interactions and overall survival analysis

Graphpad Prism 9 was used for statistical analysis. Proliferation, tumor growth rate, cell cycle, LysoTracker imaging, apoptosis, necrosis, and Cyto-ID assays were analyzed using 2-way ANOVA tests. The SynergyFinder R package was used to determine Bliss scores. A score >10 indicates a synergistic interaction, 10 to -10 indicates additivity and <-10 indicates antagonistic interaction (31). Unpaired t-tests were used to analyze the basal and TFEB siRNA invasion and migration assays, tumor necrosis, CD68 expression. One-way ANOVA was used to analyze migration and invasion assays with drug treatment. *P*-values of <0.05 were considered significant. Overall survival (OS) data is presented as Kaplan-Meier plots and were generated through Kaplan-Meier plotter (https://kmplot.com/analysis/) (23) by selecting breast cancer RNA-seq datasets and querying for *TFEB* gene and applying the filters for OS, ER+ and lymph node positive that resulted in 914 samples.

## RESULTS

### Metastatic potential of breast cancer cells is increased with abemaciclib resistance

PyMT-B6-abemaR, PyMT-Bo1-abemaR were derived from MMTV-PyMT parental cells, PyMT-B6 and PyMT-Bo1, respectively, by culturing cells in 1μM abemaciclib over 6 months (**Fig. 1A**). Basal growth rate of resistant cells remained comparable to parental sensitive cells (Supplementary Fig. S1A and B). To determine cellular sensitivity to abemaciclib, we measured cell proliferation with vehicle or increasing concentrations of abemaciclib over 72 h (**Fig. 1B and C**). PyMT-B6-abemaR and PyMT-Bo1-abemaR cells showed significant (p<0.05) decrease in sensitivity to abemaciclib at 250-2000nM compared to vehicle treated cells. The aforementioned method was also used to generate abemaciclib resistant LCC1 cells, which are MCF7 derivates that are not sensitive to estrogen (24). Similarly, LCC1-abemaR cells showed a significant decrease in sensitivity to abemaciclib at 250-2000nM (**Fig. 1D**). Moreover, PyMT-B6-abemaR and PyMT-Bo1-abemaR showed significant cross-resistance to both palbociclib and ribociclib (Supplementary Fig. 1SC-F). PyMT-B6-abemaR and PyMT-Bo1-abemaR cells failed to arrest in G1 phase of the cell cycle while sensitive PyMT-B6 and PyMT-Bo1 cells showed an increase in G1 cell cycle arrest as expected (**Fig. 1E** and **F**). Both resistant and parental cells are ER+ and are commonly used as models of luminal B breast cancer, as they also express low levels of progesterone receptor (PR) and human Epidermal Growth Factor Receptor 2 (HER2/ERBB2) (Supplementary Fig. S1G) (17). Abemaciclib targets CDK4/6 that complexes with cyclin D1 to phosphorylate and inactivate RB1 (25). Protein levels of CDK4, CDK6 and cyclin D1 were notably increased in resistant cells while phospho-RB1 (S780) levels remained unchanged compared to sensitive cells (**Fig. 1G**). Since both PyMT-B6 and PyMT-Bo1 cells are highly metastatic (17), we tested whether resistance to abemaciclib altered their metastatic potential using *in vitro* migration and invasion assays. Interestingly, both PyMT-B6-abemaR and PyMT-Bo1 cell lines were significantly more migratory and invasive that their respective parental sensitive cells (**Fig. 2A** and **B**). A significant increase in migration was also observed in LCC1-abemaR cells relative to parental LCC1 cells (Supplementary Fig. S1H). Moreover, following pathological evaluation of primary tumors following orthotopic injection of sensitive or resistant PyMT-B6 and PyMT-Bo1 cells showed significant increase in necrotic tissue and immune cell infiltration (**Fig. 2C-E**) while basal growth rates of the sensitive and resistant tumors were comparable (**Fig 2G** and **H**). Together, these data show that breast cancer cells that acquire resistance to abemaciclib also gain increased metastatic potential and aggression.

**Fig. 1.**
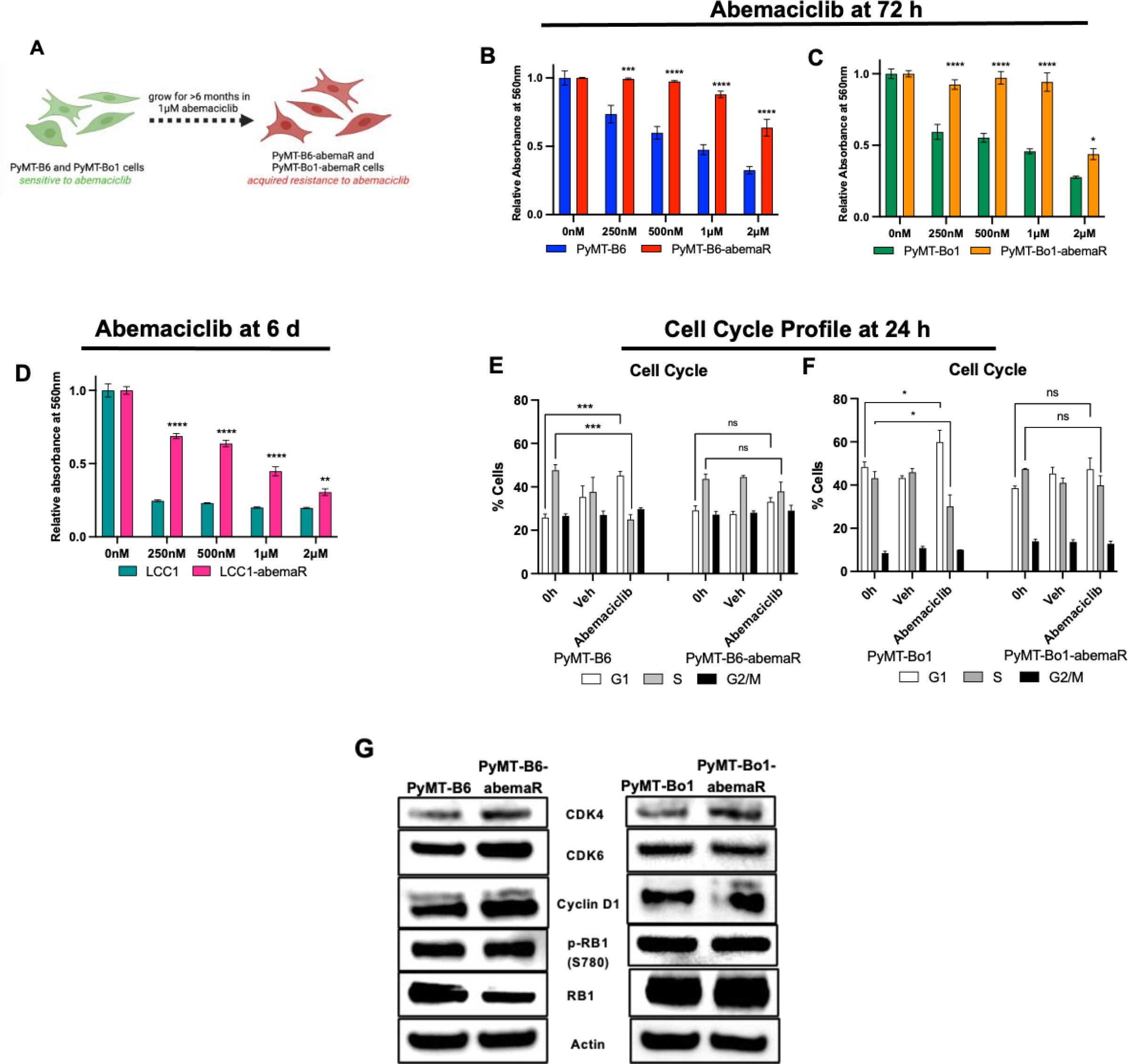
Development of abemaciclib resistant breast cancer cells. (A) Resistant cells were generated by growing sensitive cells in 1μM abemaciclib for 6 months. Cell growth rates were determined using crystal violet assays with relative cell number corresponding to absorbance determined at 560 nm, y-axes, in response to 0-2μM abemaciclib, x-axes, in (B) PyMT-B6 versus PyMT-B6-abemaR, (C) PyMT-Bo1 versus PyMT-abemaR at 72 h, and (D) LCC1 versus LCC1-abemaR cells at 6 days. *p<0.5; **p<0.01, ***p<0.001; ****p<0.0001 by two-way ANOVA. Cell cycle profile of (E) PyMT-B6 and PyMT-B6-abemaR cells, and (F) PyMT-Bo1 and PyMT-Bo1-abemaR cells after 24 h of vehicle or 1μM abemaciclib treatment. *p<0.5; ***p<0.001; ns, not significant by two-way ANOVA. (G) Western blot analysis of key CDK4/6 pathway proteins from whole cell lysates in sensitive and resistant pairs.

**Fig. 2.**
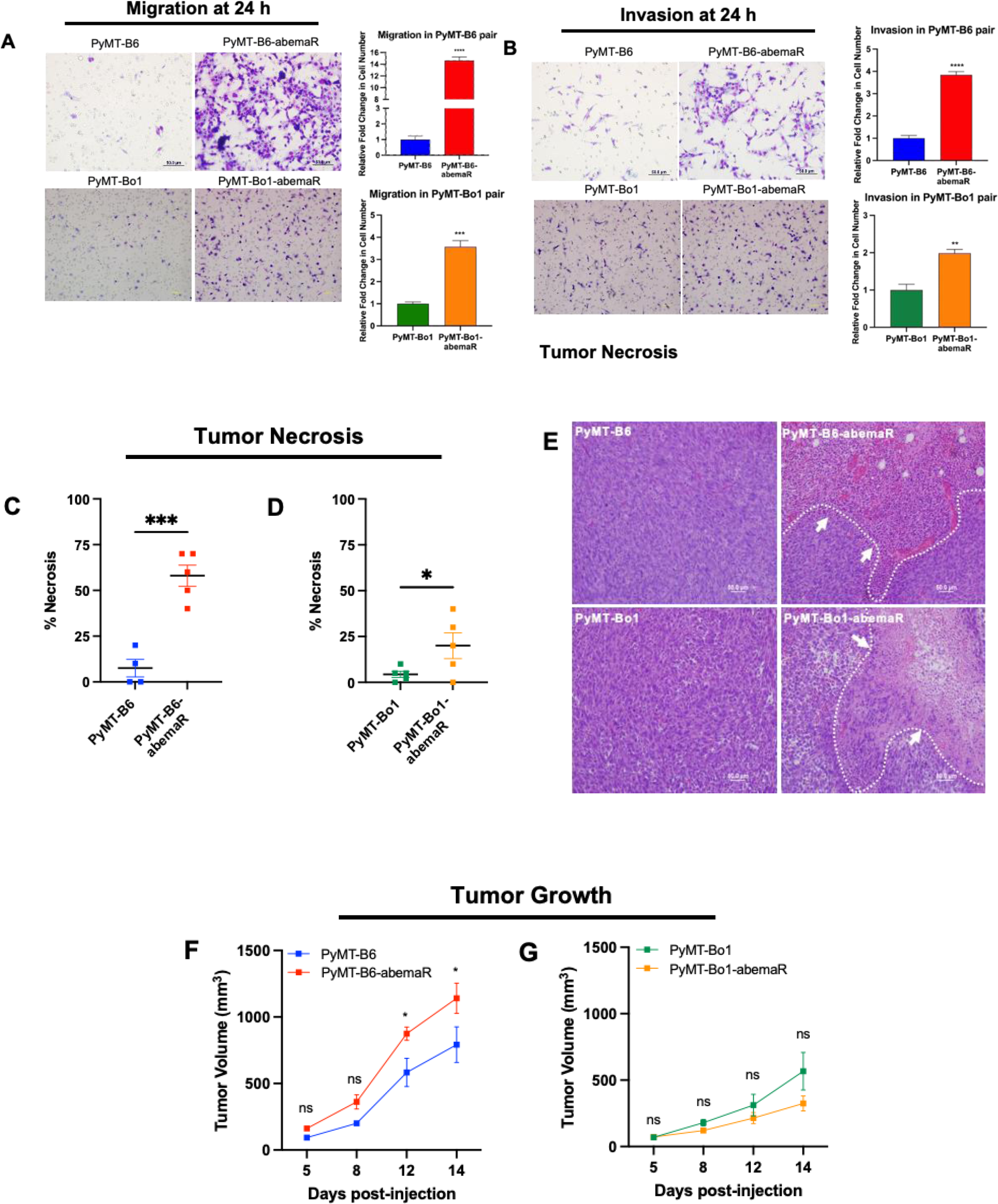
Metastatic potential is increased in abemaciclib resistant cells. PyMT-B6 and PyMT-Bo1 sensitive cells were compared to their respective resistant derivates, PyMT-B6-abemaR and PyMT-Bo1-abemaR, using transwell inserts for (A) migration and (B) invasion assays. *Right*, representative bright field images at 10× are shown. Scale bars represent 50μm for PyMT-B6 and PyMT-B6-abemaR images and 20μM for PyMT-Bo1 and PyMT-Bo1-abemaR images. *Left*, Graphs show relative total number of cells from 4 sections of transwell membranes. **p<0.01, ***p<0.001; ****p<0.0001 by Student’s t test. (C-D) Graphs show percent necrotic tissue of primary tumors formed in C57BL/6 mice following orthotopic mammary gland injections with sensitive or resistant cells. *p<0.5; ***p<0.001 by Student’s t test. (E) Representative 20× bright field images of tumor slides with H&E staining. White arrows and borders indicate necrosis. Scale bars represent 50μm. (F-G) Growth rate of tumors formed from sensitive and resistant PyMT-B6 and PyMT-Bo1 cells. *p<0.5; ns, not significant by multiple unpaired t tests.

### Tumorigenic lysosomal proteins are increased in abemaciclib resistant cells

To evaluate proteomic changes that confer resistance to abemaciclib, we conducted mass spectrometry based global proteomics analyses with sensitive PyMT-B6 and PyMT-Bo1 and their respective PyMT-B6-abemaR and PyMT-Bo1-abemaR cells. In total, >5400 proteins were identified and quantified across samples. Among them, hundreds of proteins were significantly changed between sensitive and resistant cells. Of note, very striking changes were observed for many lysosomal proteins (**Fig. 3A** and **B**, Supplementary Table S1 and S2), especially those with tumorigenic qualities such as cathepsin A, D, Z (CTSA, CTSD, CTSZ) and CD68 (lysosomal/endosomal-associated membrane glycoprotein LAMP4) (26,27). CTSD and CTSA (**Fig. 3C**) and flow cytometry for CD68 (**Fig. 3D** and **E**) confirmed increased levels of these proteins as detected in the proteomics analysis in resistant cells compared with sensitive cells. An increase in lysosomal proteins CTSD and CTSA is also observed in LCC1-abemaR cells compared to sensitive parental cells (Fig. **3F**). Previously, it was shown that abemaciclib treatment induced an increase in lysosome-based vacuole formation in treatment-naïve lung cancer cells (8). To determine if abemaciclib induced lysosomal proteins in sensitive breast cancer cells, we treated human MCF7 cells with 1 mM abemaciclib for 72 and performed quantitative proteomics (**Fig. 3G**). Indeed, up to 3% of all significantly changed proteins in abemaciclib MCF7 cells were lysosomal proteins, including CSTD and CTSA (Supplementary Table S3). Together, these data show that lysosomal proteins are remarkably increased with abemaciclib treatment and abemaciclib resistant cells retain tumorigenic lysosomal proteins.

**Fig. 3.**
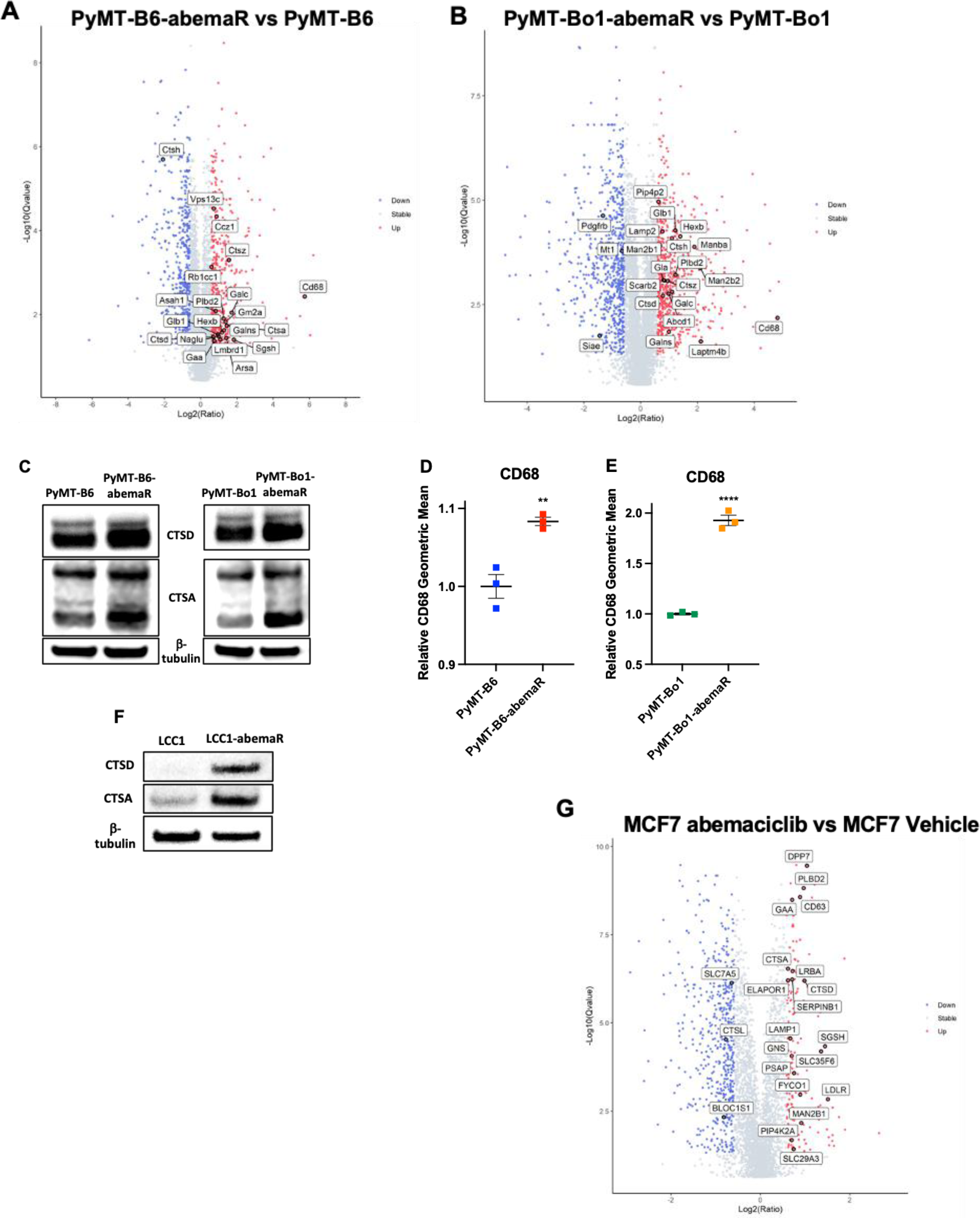
Lysosomal proteins are altered in abemaciclib resistant breast cancer cells. Volcano plots showing significantly changed proteins from mass spectrometry (MS)-based proteomics in sensitive and resistant (A) PyMT-B6 cells, and (B) PyMT-Bo1 cells. (C) Western blot analysis shows increased levels of lysosomal proteins CTSD and CTSA; β-tubulin was used as loading control. (D) Graphs show relative levels of CD68 protein captured by flow cytometry in (D) PyMT-B6 versus PyMT-B6-abemaR cells, and (E) PyMT-Bo1 versus PyMT-Bo1-abemaR cells. **p<0.01 and ****p<0.001 by Student’s t test. (F) Western blot analysis shows increased levels of CTSD and CTSA proteins in LCC1-abemaR compared to LCC1 cells. (G) Volcano plots highlighting lysosomal proteins from MS proteomics analysis of MCF7 cells after 72 h treatment with vehicle or 1μM abemaciclib.

### Lysosomal destabilizers re-sensitize resistant cells to CDK4/6 inhibitors and reduce metastatic potential

To evaluate the role of lysosomes in antiproliferative effects of abemaciclib, we treated sensitive and resistant cells with known destabilizers of lysosomal functions such as bafilomycin A1 (Baf), ammonium chloride (NH_4_Cl) and hydroxychloroquine (HCQ). Baf is a V-ATPase inhibitor that prevents lysosomal acidification and autophagosome-lysosome fusion (28), NH_4_Cl and HCQ are lysosomotropic agents that become protonated and trapped within lysosomes (29). HCQ is a clinical grade anti-malarial drug that is also used as an anticancer drug for its ability to disrupt lysosome-autophagosome fusion (30). Since Baf is highly toxic to all cells within 24 h, dose response with increasing concentrations of Baf was done within 15 h (Supplementary Fig. S2A and B) while dose responses with NH_4_Cl and HCQ were done for 72 h. Abemaciclib resistant cells were more sensitive to Baf compared to sensitive cells in both PyMT-B6 and PyMT-Bo1 pairs. On the other hand, dose-responses with NH_4_Cl and HCQ showed increased sensitivity in abemaciclib sensitive cells compared to resistant cells (Supplementary Fig. S2C-F). Next, we combined a single dose of Baf or HCQ, based on efficacy as single agent, with increasing doses of abemaciclib (Supplementary Fig. S3A-D, **Fig. 4A-D**). To determine synergistic interaction between abemaciclib and HCQ, we calculated Bliss synergy scores (31) and showed that these drugs interact synergistically, with scores >10, in both sensitive and resistant PyMT-B6 and PyMT-Bo1 cells (Supplementary Fig. S3I-J). Notably, abemaciclib and HCQ were more synergistic in resistant cells at doses of abemaciclib >1 μM in both cell lines. Resistant LCC1-abemaR cells are also are re-sensitized to abemaciclib with the addition of HCQ (**Fig. 4E** and **F**). PyMT-Bo1 cells were also made resistant to palbociclib in the same manner as the abemaciclib resistant cells (Supplementary Fig. S4A). Both palbociclib sensitive and resistant PyMT-Bo1 cells were further sensitized to palbociclib with the addition of 5μM HCQ (Supplementary Fig. S4B and C). Additionally, sensitive PyMT-Bo1 cells and cells made resistant to ribociclib (Supplementary Fig. S4D) experienced reduced proliferation when abemaciclib was combined with HCQ (Supplementary Fig. S4E and F). Since PyMT-B6 and PyMT-Bo1 cells are metastatic breast cancer cell models (17), we asked if resistance to abemaciclib changes their migratory or invasive potential. Results from *in vitro* migration (Supplementary Fig. S5A and B) showed that while abemaciclib alone inhibited cell migration in PyMT-B6, PyMT-B6-abemaR and PyMT-Bo1-abemaR cells, the combination of HCQ and abemaciclib is significantly more effective in inhibiting migration in these cells. Though, abemaciclib alone or the combination of HCQ and abemaciclib did not change migration in PyMT-Bo1 cells. Results from invasion (**Fig. 4G** and **H**) assays showed that combination of abemaciclib and HCQ significantly inhibited invasion in both sensitive PyMT-B6 and resistant PyMT-B6-abemaR cells as well as PyMT-Bo1-abemaR cells compared to vehicle alone. Notably, in PyMT-Bo1 cells, neither treatment with monotherapies nor the combination affected invasive potential of the cells. Thus, inhibition of lysosomal function inhibited migration and invasion potential in both abemaciclib sensitive and resistant variants of PyMT-B6 cells. However, in PyMT-Bo1 cells, migration and invasion was significantly inhibited only in the resistant PyMT-Bo1-abemaR cells. Together, these data highlight an essential role of lysosomes in promoting migratory and invasive potential in metastatic breast cancer cells. Additionally, these data also show that targeting lysosomes can inhibit migratory and invasive potential in abemaciclib resistant cells.

**Figure 4.**
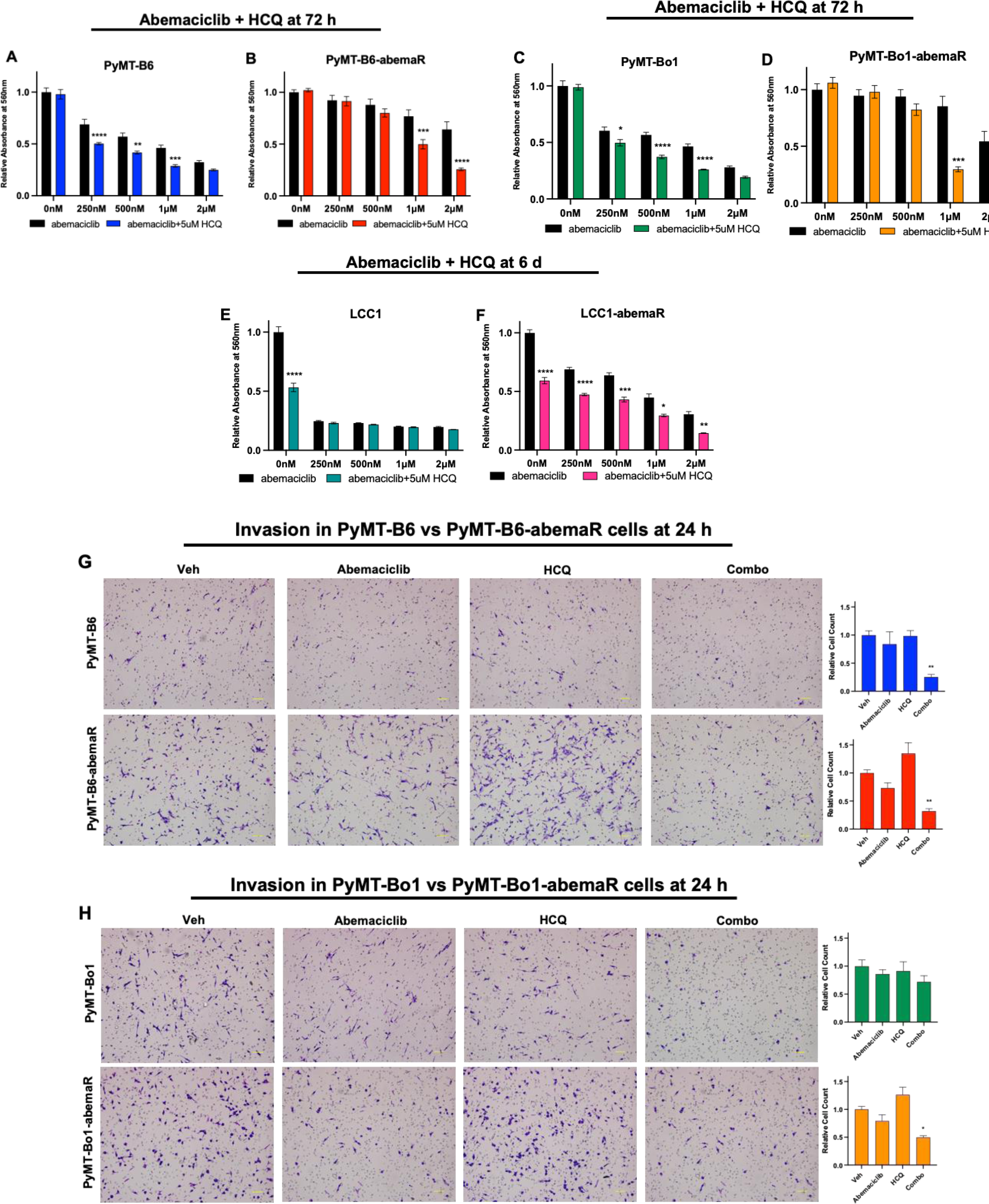
Combination of abemaciclib and HCQ inhibits cell proliferation and invasion in resistant cells. (A-D) Cell growth rates were determined using crystal violet assays with relative absorbance at 560 nm corresponding to cell number in response to 0-2μM abemaciclib+ 5μM HCQ for 72 h. (E,F) Cell growth in LCC1 cells to 0-2μM abemaciclib+5μM HCQ for 6 days. *p<0.5; **p<0.01, ***p<0.001; ****p<0.0001 by two-way ANOVA. (G,H) *In vitro* invasion assay of PyMT-B6 or PyMT-Bo1 (top panels) and resistant PyMT-B6-abemaR and PyMT-Bo1-abemaR (bottom panels) cells with vehicle, 500nM abemaciclib, 5μM HCQ, or the combination treatment for 24 h. Trapped cells on membranes were stained with H&E and separate fields of views at 10× bright field images were counted and the total number of cells was averaged in the corresponding graphs. Scale bars represent 100μm. *p<0.05 and **p<0.01 by one-way ANOVA.

### Abemaciclib resistant cells are more susceptible to lysosomal membrane permeabilization than sensitive cells

Lysosomes are acidic single membrane organelles that contain >50 different acid hydrolases. Induction of lysosomal membrane permeabilization (LMP) leads to lysosome dependent cell death (LDCD) (32,33). To asses LMP in abemaciclib sensitive and resistant cells, we incubated PyMT-B6, PyMT-B6-abemaR, PyMT-Bo1 and PyMT-Bo1-abemaR cells with LysoTracker Green and evaluated lysosomal staining with live confocal microscopy. Treatment with 1μM abemaciclib induced robust accumulation of LysoTracker within lysosomes compared to vehicle control. Treatment with 5μM HCQ did not show a marked difference in LysoTracker staining compared to control. However, treatment with combination of abemaciclib and HCQ diminished LysoTracker accumulation (**Fig. 5A** and **B**). Since LMP can induce cell death through a variety of pathways, we compared the percentage of cells undergoing apoptosis using annexin V assays (**Fig. 6A** and **B**) in PyMT-B6 and PyMT-Bo1 abemaciclib sensitive and resistant pairs under the different treatment conditions. We found that treatment with abemaciclib alone or the combination of abemaciclib and HCQ induced a significant but modest increase (∼10%) in apoptosis in sensitive PyMT-B6 cells compared with vehicle treatment. In PyMT-B6-abemaR cells, a significant but modest (∼5%) increase in apoptotic cells were present only with the combination treatment compared with vehicle. In PyMT-Bo1 cells, abemaciclib alone or the combination of abemaciclib and HCQ induced a significant and robust increase (40%) in apoptotic cells. While abemaciclib alone or in combination with HCQ induced a relatively lesser effect (20%) compared to vehicle in PyMT-Bo1-abemaR cells. Treatment with HCQ alone did not show an increase in apoptotic cells in either of the four cell lines. Next, we determined autophagy induction by measuring accumulation of autophagosomes in both sensitive and resistant cells (**Fig. 6C** and **D**). Following treatment with either abemaciclib alone or the combination of abemaciclib and HCQ, there was approximately a 4-fold and 2-fold significant induction of autophagosomes in PyMT-B6 cells and PyMT-B6-abemaR cells, respectively. In PyMT-Bo1 cells, induction of autophagosomes was 2-fold with abemaciclib alone or in combination with HCQ. While no significant differences were seen with any treatment conditions in PyMT-Bo1-abemaR cells. Serum deprived starvation was used as positive control in detecting autophagosomes. Interestingly, we noticed a varied response between sensitive and resistant cells to starvation, although in all cells, starvation induced increased accumulation of autophagosomes. To further confirm the induction of cell death pathways, we conducted Western blotting with known markers apoptosis and autophagy (**Fig. 6E**). In both PyMT-B6 and PyMT-Bo1 sensitive cells, abemaciclib alone, or more markedly, abemaciclib and HCQ treatment induced an increase in cleaved PARP1 (poly(ADP-ribosyl)transferase), a known marker of apoptosis (34). In contrast, cleaved PARP1 was not increased in PyMT-B6-abemaR and PyMT-Bo1-abemaR cells with abemaciclib alone or in combination with HCQ compared to vehicle. To measure autophagic flux, we examined protein levels of p62 (SQSTM1), an autophagosome cargo marker that decreases in active autolysosomes, and LC3B-II, a marker that increases with formation and enlargement of autophagosome (35). Abemaciclib alone decreased p62 levels and increased LC3B-II compared to vehicle in sensitive PyMT-B6 cells only. HCQ alone induced a modest increase in LC3B-II without a concomitant decrease in p62 compared to vehicle in all cell lines suggesting incomplete autophagy. Combination of HCQ and abemaciclib increased LC3B-II levels with simultaneous decrease in p62 in sensitive PyMT-B6 and PyMT-Bo1 cells but not in respective resistant cells. Considering the relatively subdued induction of apoptosis or autophagy in resistant cells, we examined the protein levels of BCL2 (B cell lymphoma 2), a well-characterized inhibitor of multiple death pathways, including apoptosis, necrosis, and autophagy (36,37). Interestingly, BCL2 levels were increased in resistant cells compared to sensitive cells in both PyMT-B6 and PyMT-Bo1 cell lines (**Fig. 6F**) and remained detectable following treatment with HCQ, abemaciclib or the combination in resistant cells. Paclitaxel (50 nM) was used as a control due to its inhibitory effect on BCL2 expression (38). Furthermore, BCL2 protein was markedly increased in LCC1-abemaR cells compared to LCC1 cells (**Fig. 6G**). Since increased levels of BCL2 protein were detected in abemaciclib resistant cells, we tested the efficacy of BCL2 inhibitors, venetoclax (VNX) (**Fig. 6H**) and obatoclax (OBT) (Supplementary Fig. S6A) in PyMT-Bo1 and PyMT-Bo1-abemaR cells. Based on these dose-response studies, we used the IC_50_ concentrations for VNX (7μM) and OBT (15nM), in PyMT-Bo1 cells, to test the efficacy of triple combination of HCQ, abemaciclib and a BCL2 inhibitor in abemaciclib sensitive and resistant cells. Combination of HCQ, abemaciclib and VNX had the same inhibitory effect as HCQ and abemaciclib in PyMT-Bo1 cells (**Fig. 6I**). However, the combination of HCQ, abemaciclib and VNX significantly decreased cell proliferation compared to the combination of HCQ and abemaciclib in PyMT-Bo1-abemaR cells (**Fig. 6J**). Combination of HCQ, abemaciclib and OBT significantly decreased cell proliferation compared to the combination of HCQ and abemaciclib in both PyMT-Bo1 and PyMT-Bo1-abemaR cells (Supplementary Fig. S6B and C). VNX specifically inhibits BCL2 while OBT is a pan-BCL2 inhibitor and can inhibit multiple members of the BCL2 family, including BCL2L1 (BCLxL) and MCL1 (39). Together, these data show that increased levels of BCL2 contributes to acquired resistance to abemaciclib, and supports the use of anti-BCL2 drugs as a therapeutic option in acquired resistance to CDK4/6i.

**Figure 5.**
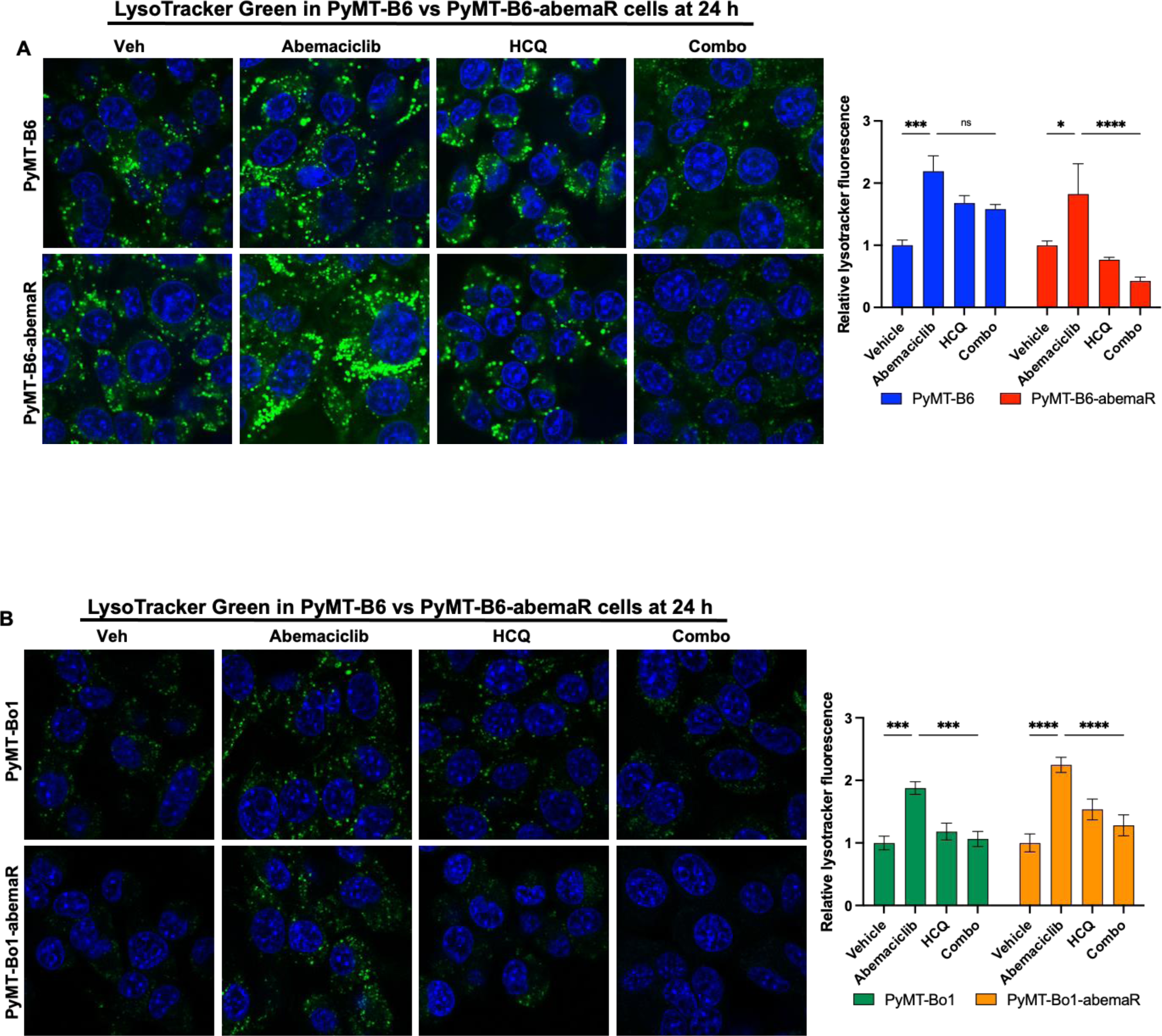
Combination of abemaciclib and HCQ induces lysosomal membrane permeability (LMP) Lysosomes were imaged with live microscopy following staining cells with LysoTracker Green (green punctates) after 24 h treatment with vehicle, 1μM abemaciclib, 5μM HCQ or the combination in (A) *left*, representative images for PyMT-B6 and PyMT-B6-abemaR cells; *right*, graph showing changes in relative lysotracker fluorescence, and (B) *left*, PyMT-Bo1 and PyMT-Bo1-abemaR cells; *right*, graph showing changes in relative lysotracker fluorescence. *p<0.05, ***p<0.001, ****p<0.0001, ns, not significant, by one-way ANOVA. Nuclei were stained with Hoechst 33342 (blue). All images were obtained at 60× magnification.

**Figure 6.**
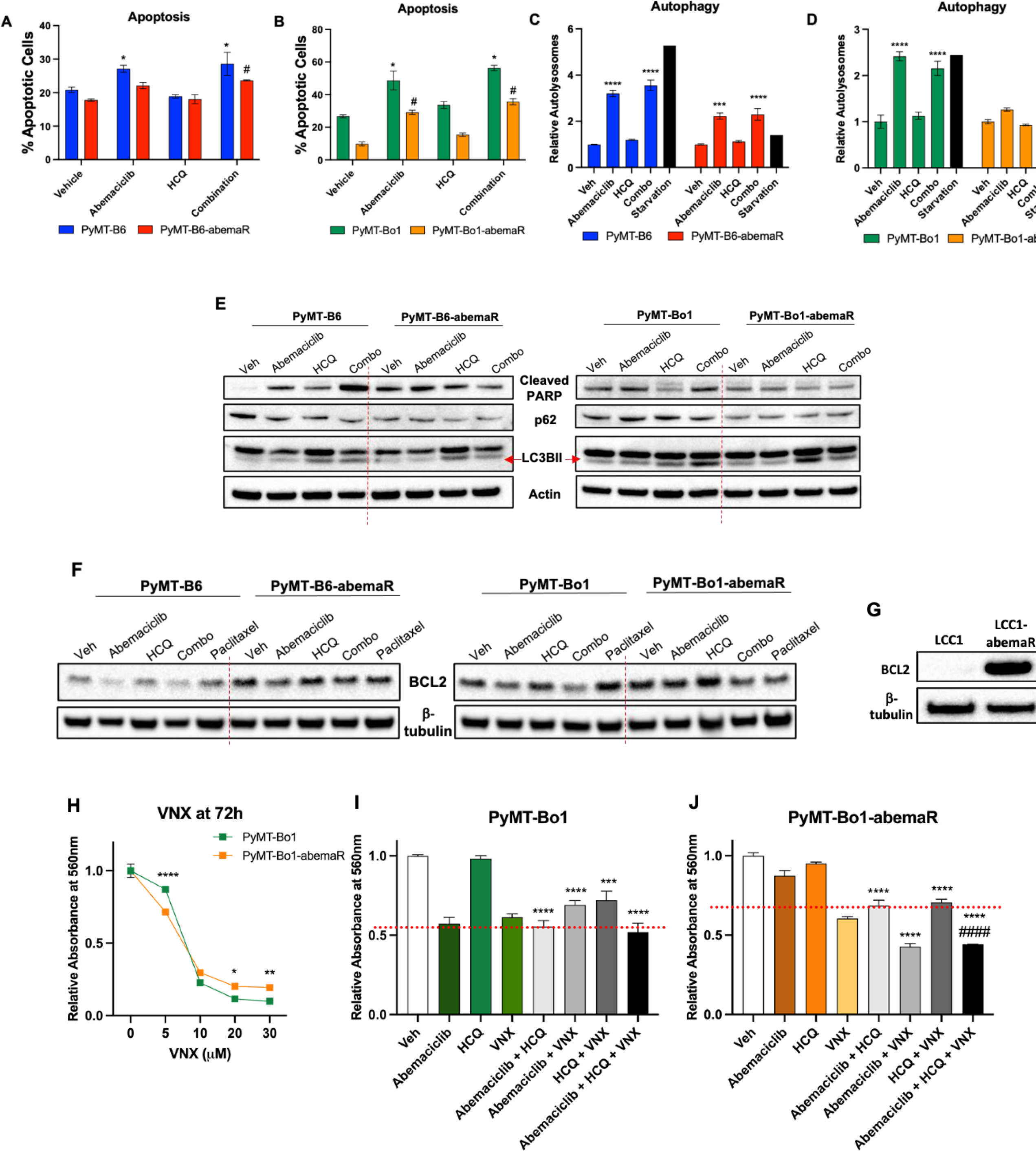
Cell death via apoptosis and autophagy are reduced in abemaciclib resistant cells compared to sensitive cells. (A,B) Percent apoptosis levels in PyMT-B6 and PyMT-B6-abemaR cells following treatment with vehicle, 1μM abemaciclib, 5μM HCQ or the combination for 72 h detected by flow cytometry using annexin V assays. (C,D) Relative autolysosomes detected by flow cytometry using Cyto-ID assays in PyMT-B6, PyMT-abemaR, PyMT-Bo1 and PyMT-Bo1-abemaR cells after treatment with vehicle, 1μM abemaciclib, 5μM HCQ or the combination for 72 h. *#p<0.5; ***p<0.001; ****p<0.0001 compared to respective vehicle by two-way ANOVA. (E) Western blots show induction of apoptosis (cleaved PARP1) and autophagy (p62, LC3BII) markers after treatment with vehicle, 1μM abemaciclib, 5μM HCQ or the combination for 72 h. (F) Western blots show levels of BCL2 proteins in cells under specified treatment conditions: vehicle, 1μM abemaciclib, 5μM HCQ or the combination for 72 h; 50 nM paclitaxel was used as a control for inhibiting BCL2. (G) Western blot shows increased BCL2 protein levels in LCC1-abemaR compared to LCC1 cells under basal conditions. β-tubulin was used as a loading control. (H) Cell growth in response to 5-30 μM BCL2 inhibitor venetoclax (VNX) in PyMT-Bo1 and PyMT-Bo1-abemaR cells at 72 h. (I, J) Cell growth in PyMT-Bo1 and PyMT-Bo1-abemaR cells treated with vehicle, 500nM abemaciclib, 5μM HCQ, 7 μM VNX or the dual and triple combinations for 72 h. Red dashed line marks the inhibitory effect of the combination of HCQ+abemaciclib in respective cell lines. ***p<0.05 and ****p<0.01 compared to vehicle (Veh); ####p<0.0001 compared to HCQ+abemaciclib by one-way ANOVA.

To determine whether combination of abemaciclib and HCQ affected cell cycle, we analyzed cell cycle profiles in sensitive and resistant pairs of PyMT-B6 and PyMT-Bo1 cells. As expected, treatment with abemaciclib alone induced G1 cell cycle arrest in both sensitive cells, PyMT-B6 and PyMT-Bo1, however, combination of abemaciclib and HCQ did not further augment G1 cell cycle arrest in sensitive cells (Supplementary Fig. 76A-D). In resistant PyMT-B6-abemaR and PyMT-Bo1-abemaR cells, neither abemaciclib alone or the combination of abemaciclib and HCQ affected cell cycle profiles compared to vehicle control. Collectively, these data show that combination of abemaciclib and HCQ induce LMP in abemaciclib resistant cells, however, apoptosis, autophagy or cell cycle arrest are not robustly induced as compared to sensitive cells.

### Genetic alteration of *TFEB* is associated with tumorigenesis in ER+ breast cancer

TFEB is a master regulator of a number of lysosomal and autophagy proteins. Since lysosomal proteins were deregulated in abemaciclib resistant cells, we analyzed TFEB protein levels in abemaciclib sensitive and resistant cells. Western blot analysis showed increased TFEB protein levels in PyMT-B6-abemaR and PyMT-Bo1-abemaR cells compared to their sensitive parental cells (**Fig. 7A**). Knockdown of TFEB with siRNA (**Fig. 7A**) showed no change in abemaciclib mediated inhibition of cell proliferation with vehicle or abemaciclib treatment at 72 h (Supplementary Fig. 8A-D) but showed a significant decrease in invasive potential of PyMT-B6-abemaR and PyMT-Bo1-abemaR cells (**Fig. 7B** and **C**). TFEB has previously been reported to affect invasion but not proliferation in human squamous carcinoma cells (40). Next, we analyzed 7,298 invasive breast cancer patient sample data curated by cBioportal, and found amplifications, deletions and other mutations in 2% of all cases (**Fig. 7D**). Furthermore, analysis of overall survival (OS) in ER+ breast cancer patients showed that increased TFEB expression correlated with lower OS (HR = 2.62 [1.35-5.07]; *p*=0.003) (**Fig. 7E**). Together, these data show that increased or genetically altered *TFEB* expression in breast cancer correlated with increased tumorigenesis in a subset of breast cancer.

**Figure 7.**
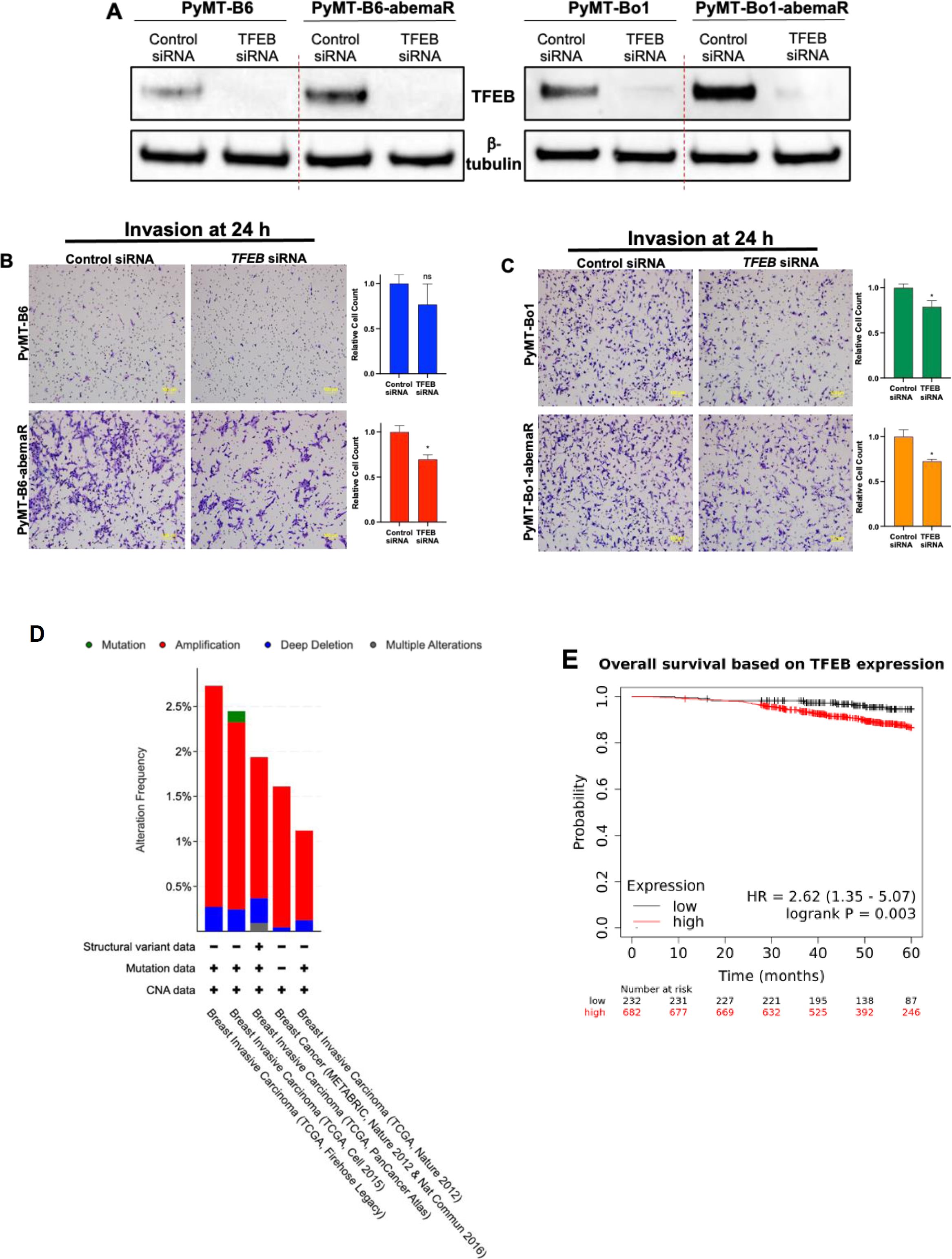
Genetic alteration of TFEB is associated with tumorigenesis in ER+ breast cancer. (A) Western blot of TFEB in PyMT cells after 72h transfection with 20nM control of TFEB siRNA. (B,C) Invasion of sensitive (top) and resistant (bottom) PyMT cells after 24h transfection with 20nM control or TFEB siRNA. Cells from four images were counted and the total number of cells was averaged in the corresponding graphs. Scale bars represent 100μm. *p<0.5; ns, not significant by Student’s t test. (D) *TFEB* mutational status in breast tumors of metastatic patients acquired through Cbioportal. (E) Overall survival curves of node-positive ER+ breast cancer patients stratified by *TFEB* gene expression in primary tumors.

## DISCUSSION

CDK4/6i are used in combination with antiestrogens as the first-line treatment of ER+ metastatic breast cancer, and abemaciclib in particular is used in the adjuvant setting for patients with early, high-risk disease (4). Although CDK4/6i successfully extend progression-free survival, resistance is a common problem, and currently, there are no FDA approved drugs to specifically treat CDK4/6i resistant breast cancer (41). To this point, research into resistance had focused on implicating growth factors or cell cycle proteins (42), however, efficacy of these agents remain clinically unconfirmed. Recently, lysosomes have been implicated in a non-apoptotic mode of cell death in response to abemaciclib and as a mode of inherent resistance to palbociclib in triple negative breast cancer (8,9). Similarly, we show that in sensitive MCF7 cell, abemaciclib induced an increase of lysosomal proteins (**Fig. 3G**; Supplementary Table S3) compared to vehicle treatment. Hence, abemaciclib triggers activation of cellular mechanisms that increases lysosomal proteins in sensitive cells. Here, we show that metastatic breast cancer cells that acquire resistance to abemaciclib attain increased metastatic potential and contain higher levels of lysosomal proteins levels, which are associated with tumorigenesis. Therefore, targeting the lysosomal pathway is a plausible therapeutic strategy in advanced ER+ metastatic breast cancer.

Lysosomes are primarily known for catabolic turnover of various macromolecules but these versatile organelles are also regulators of cell proliferation and can promote metastasis (16,43). Lysosomes are regulated by the microphthalmia/transcription factor E (MiT/TFE) family of transcription factors consisting of MITF, TFE3, TFEC, and TFEB (44). TFEB has many functions, but regulation of lysosomal-autophagy genes is one that can directly contribute to metastasis (45). In our abemaciclib resistant breast cancer models, while we do not see changes in lysosomal number with vehicle alone or abemaciclib (**Fig. 5A** and **B**), levels of a number of lysosomal proteins were significantly changed (**Fig. 3**). Further studies are needed to determine how increase in TFEB alters the lysosomal proteins in CDK4/6i resistant cells. Through analyses of publicly available human breast cancer data, we show that *TFEB* mutations, particularly amplification, are present in a subset of invasive breast cancer cases, and increased *TFEB* gene expression correlated with decreased OS in ER+ breast cancer cases (**Fig. 7D** and **E**). Evaluation of TFEB protein levels in 100 breast cancer patients with small tumors (<2 cm) showed high TFEB expression in one-fourth of the cases (46). Increased TFEB was found to be a prognostic variable and also positively correlated with increase in the autophagosome gene *LC3A* levels in this cohort. Therefore, it is possible that in a subset of breast tumors that harbor increased TFEB levels, treatment with CDK4/6i will lead to poor clinical outcome, and targeting TFEB-mediated pro-survival pathways could improve efficacy of CDK4/6i in these cases.

HCQ is widely used as an inhibitor of the autophagy-lysosomal pathway (47). HCQ is a weak base that accumulates in acidic compartments of cells and tissue. HCQ can disrupt lysosomal function, autophagy, membrane integrity as well as transcription and signaling (48). Like HCQ, abemaciclib is lysosomotropic (9) and the combination of these two drugs disrupts lysosomal integrity by inducing LMP, and downstream death, in resistant cells (**Fig. 5A** and **B**). We noted a reduction in death in resistant cells compared to sensitive cells, and found an upregulated of BCL2 in these cells (**Fig. 6**). Recently, two trials have begun to test the efficacy of HCQ or BCL2 inhibitors with CDK4/6i in ER+ breast cancer. The ABBY trial is investigating the efficacy of adding HCQ, as an autophagy inhibitor, to abemaciclib in CDK4/6 inhibitor-naïve patients to prevent survival of senescent breast cancer cells disseminated in the bone (49). The PALVEN trial is designed to test the safety of palbociclib, letrozole, and a BCL2 inhibitor, venetoclax, as adding a BCL2 inhibitor to CDK4/6i triggers higher levels of apoptosis in cancer cells (50). These studies provide evidence that adding targeted therapies, which disrupt survival signaling that inhibit cell death pathways, to CDK4/6i can increase their efficacy. Future studies will investigate the effects of combining abemaciclib and other lysosome destabilizing drugs on tumor growth and metastasis using *in vivo* models of ER+ breast cancer.

## ACKNOWLEDGEMENTS

Funding support for this study is in part from Georgetown University Medical Center. Technical services were provided by the following shared resources at Georgetown University Medical Center: Tissue Culture, Flow Cytometry, Animal Models and Mass Spectrometry and Analytical Pharmacology Shared Resource that were funded through Public Health Service award P30-CA-51008 (Lombardi Comprehensive Cancer Center Support Grant). The instrument Orbitrap Lumos Tribrid mass spectrometer was partially supported by Dekelbaum Foundation. We thank Idalia Cruz for technical assistance with the *in vivo* studies, and Dr. Susana Galli (veterinary pathologist) for pathological evaluation of the murine primary tumor samples. We also thank the Georgetown Breast Cancer Advocates (GBCA) for a patient’s perspective for the translational aspect of our study.

---

**Figure S1.**
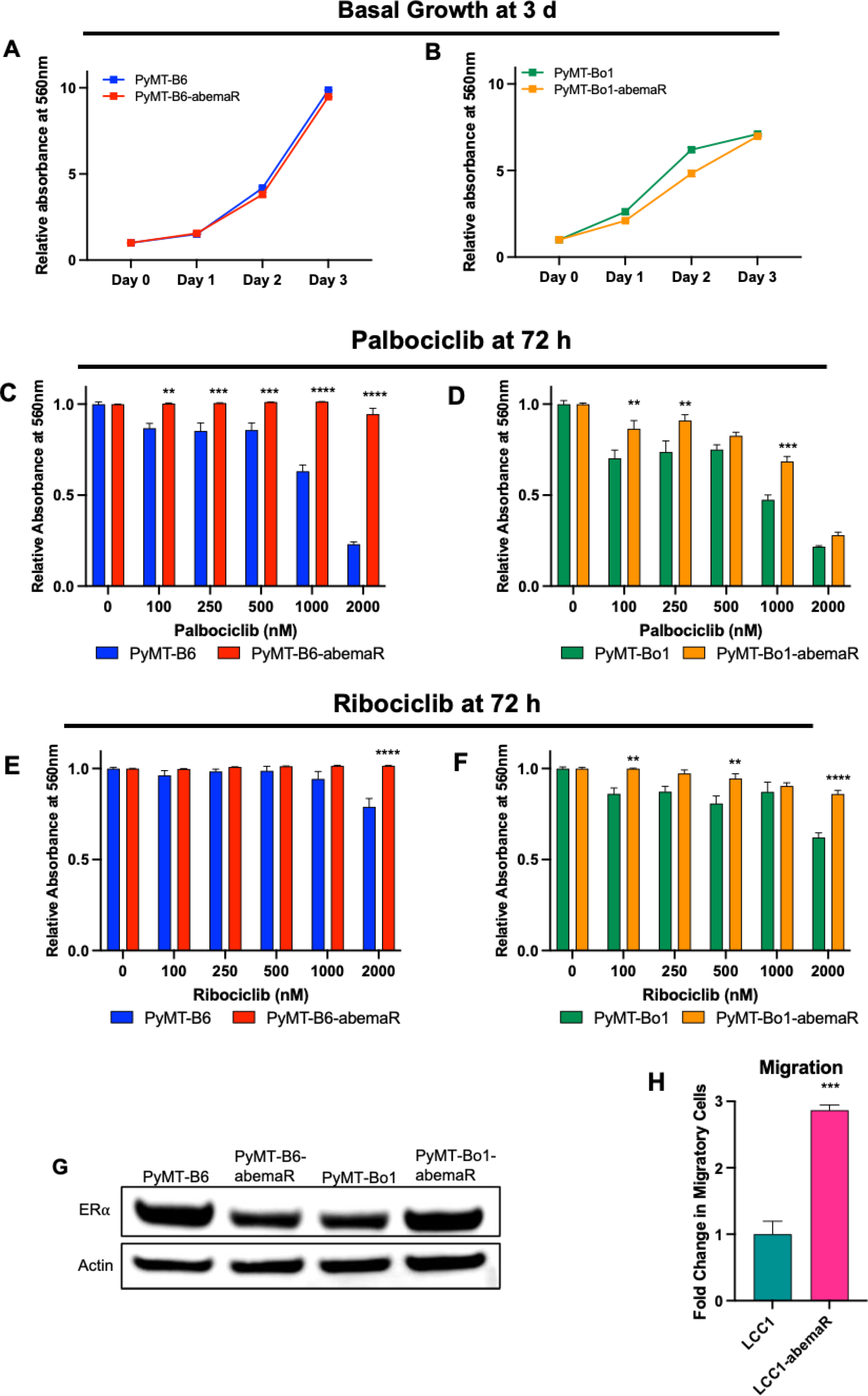
Abemaciclib resistant cells are cross-resistant to palbociclib and ribociclib. (A) Growth rates of PyMT-B6 and PyMT-B6-abemaR, and (B) PyMT-Bo1 and PyMT-Bo1-abemaR cells over 3 days; relative growth was normalized to cell number on day 0. Cell number was determined with crystal violet assays. Cell growth of PyMT-B6 versus PyMT-B6-abemaR, and PyMT-Bo1 versus PyMT-Bo1-abemaR cells after growing in indicated concentrations of (C, D) palbociclib or (E, F) ribociclib for 72h. **p<0.01, ***p<0.001; ****p<0.0001 by two-way ANOVA. (G) Western blot to show ER⍺ protein levels PyMT-B6, PyMT-B6-abemaR, PyMT-Bo1 and PyMT-Bo1-abemaR cells. (H) LCC1-abemaR cells are significantly more migratory than parental sensitive LCC1 cells; graph shows relative fold change in migratory cells in an *in vitro* migration assay 48 h assay. ***p<0.001 by Student’s t test.

**Figure S2.**
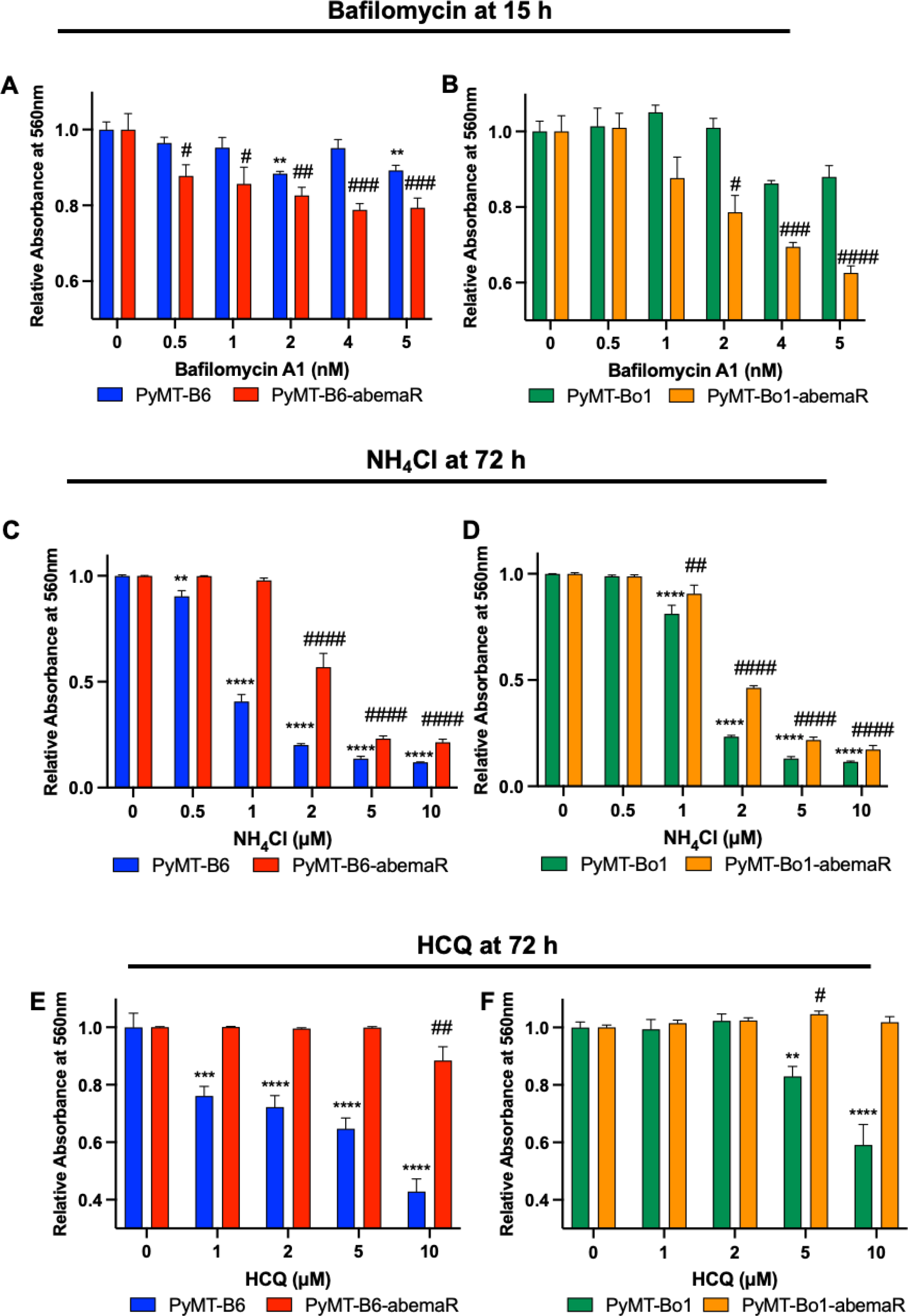
Abemaciclib resistant cells are more sensitive to bafilomycin A1 and less sensitive to NH_4_Cl and HCQ compared to sensitive cells. Cell growth was measured by crystal violet assays in response to 72 h treatment with: 0.5-5 nM bafilomycin A1 (Baf) in (A) PyMT-B6 and PyMT-B6-abemaR, (B) PyMT-Bo1 and PyMT-Bo1-abemaR; 0-10μM NH_4_Cl in (C) PyMT-B6 and PyMT-B6-abemaR, (D) PyMT-Bo1 and PyMT-Bo1-abemaR; 0-10 μM HCQ in (E) PyMT-B6 and PyMT-B6-abemaR, (F) PyMT-Bo1 and PyMT-Bo1-abemaR. *#=p<0.05, **##=p<0.01, ***###=p<0.001, ****####=p<0.0001 by two-way ANOVA compared to 0 concentration of drug for respective cell line.

**Figure S3.**
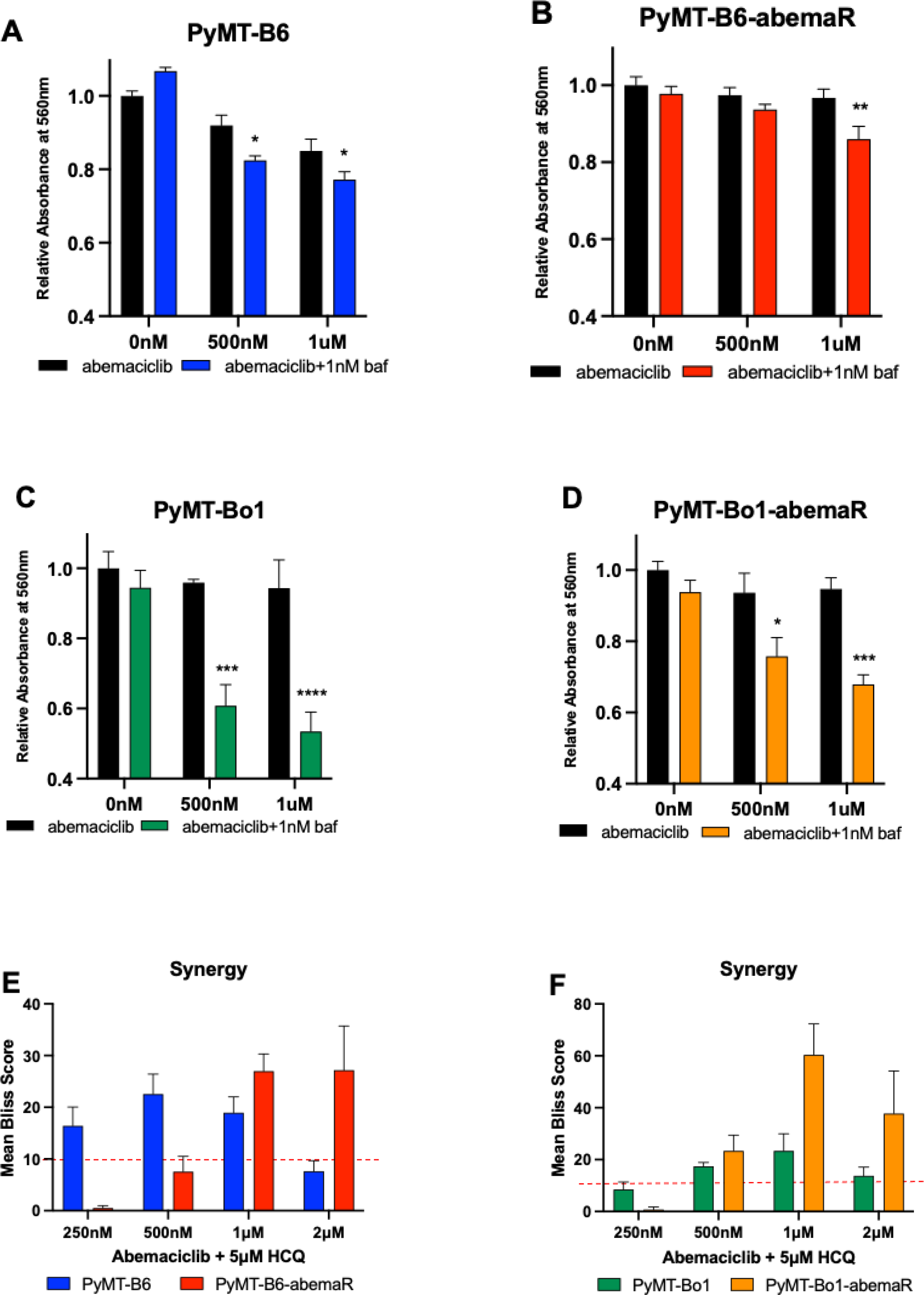
Abemaciclib and lysosomal destabilizing drugs re-sensitize resistant cells to abemaciclib. (A-D) Cell growth was determined in response to increasing doses (0-1μM) of abemaciclib in the presence of 1nM bafilomycin for 15h. *p<0.5; **p<0.01, ***p<0.001; ****p<0.0001 by two-way ANOVA. (E, F) Mean Bliss scores for indicated concentrations of abemaciclib and 5μM HCQ using the SynergyFinder R package. A score >10 (above red dashed line) indicates a synergistic interaction and 10 to -10 indicates an additive interaction between abemaciclib + 5μM HCQ.

**Figure S4.**
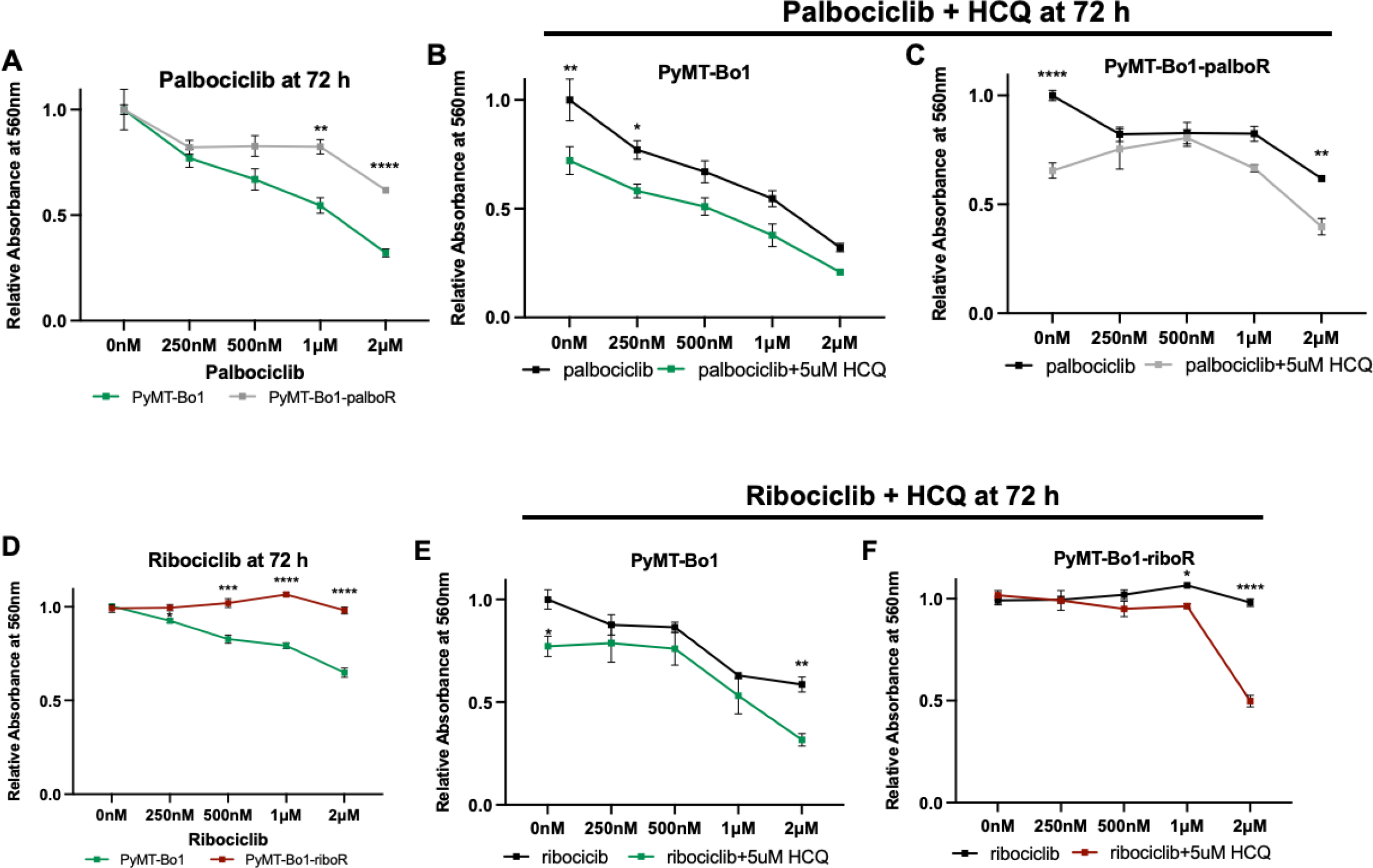
HCQ re-sensitizes palbociclib and ribociclib resistant cells to their respective CDK4/6i. (A) Cell growth in PyMT-Bo1 and PyMT-Bo-1-palboR cells in response to 0-2μM palbociclib at 72h. Cell growth in (B) PyMT-Bo1 and (C) PyMT-Bo1-palboR cells following treatment with 0-2μM palbociclib + 5μM HCQ at 72h. (D) Cell growth in PyMT-Bo1 and PyMT-Bo1-riboR cells in response to 0-2μM ribociclib at 72h. Cell growth in (E) PyMT-Bo1 and (F) PyMT-Bo1-riboR cells following treatment with 0-2μM ribociclib + 5μM HCQ at 72h. *p<0.5; **p<0.01, ***p<0.001; ****p<0.0001 by two-way ANOVA comparing cell growth between the cell lines.

**Figure S5.**
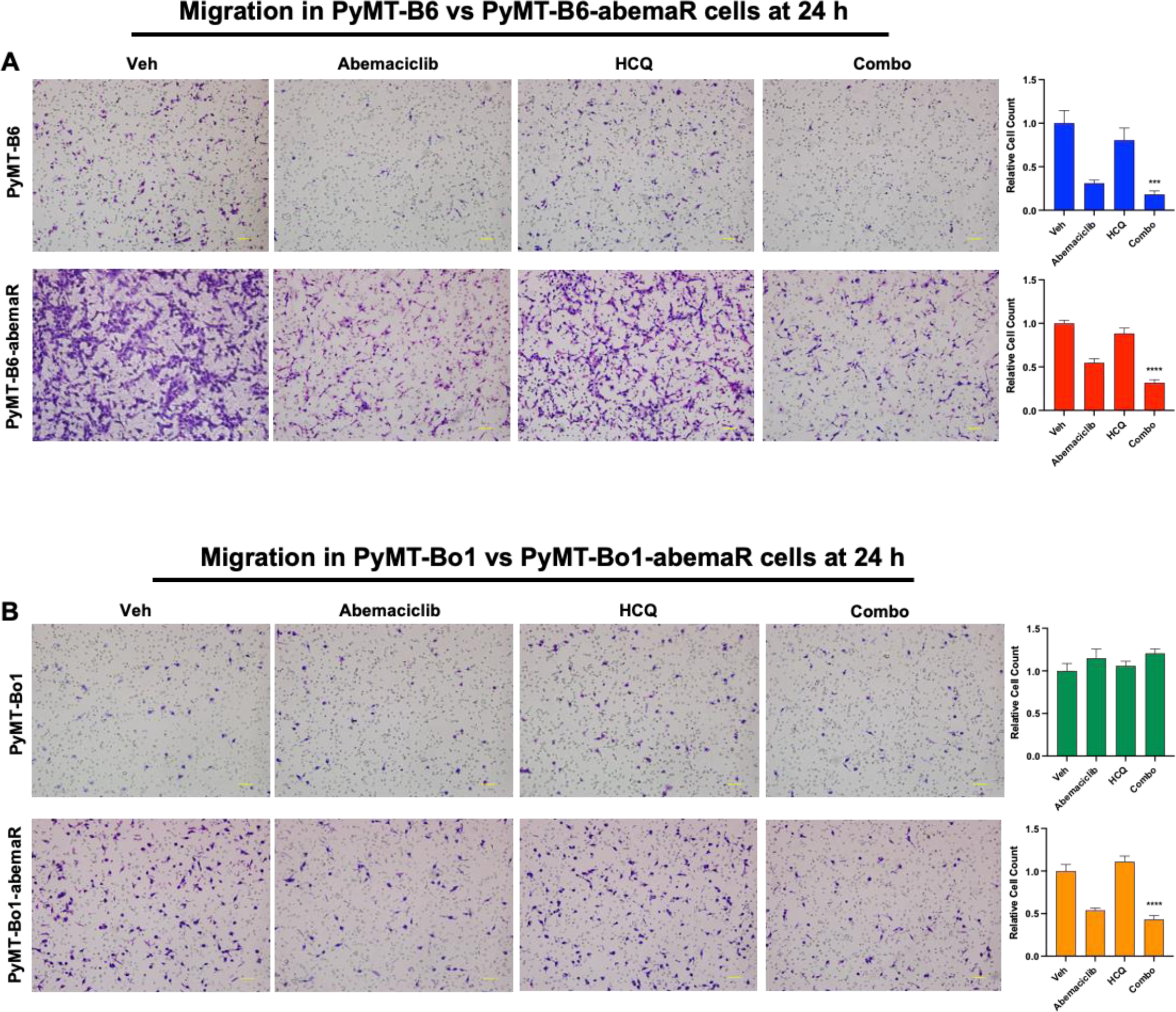
Combination of abemaciclib and HCQ inhibits cell migration in resistant cells. (A,B) *In vitro* migration assay with vehicle, 500nM abemaciclib, 5μM HCQ, or the combination treatment for 24h. *In vitro* migration assay of PyMT-B6 or PyMT-Bo1 (top panels) and resistant PyMT-B6-abemaR and PyMT-Bo1-abemaR (bottom panels) cells with vehicle, 500nM abemaciclib, 5μM HCQ, or the combination treatment for 24 h. Trapped cells on membranes were stained with H&E and separate fields of views at 10× bright field images were counted and the total number of cells was averaged in the corresponding graphs. Scale bars represent 100μm. ***p<0.05 and ****p<0.01 by one-way ANOVA.

**Figure S6.**
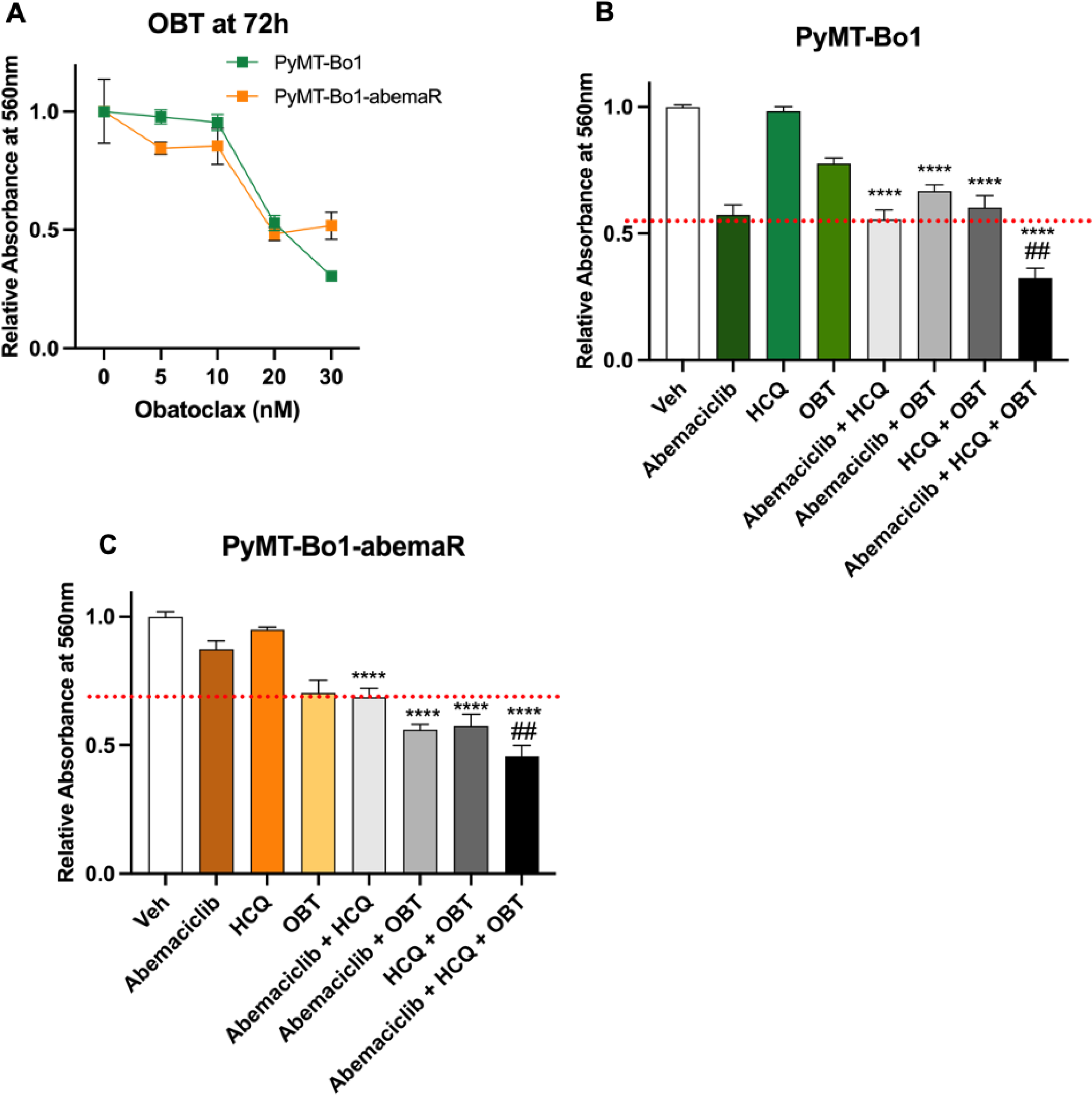
Combination of abemaciclib, HCQ and BCL2 inhibitor obatoclax inhibits cell migration in both abemaciclib sensitive and resistant cells. (A) Cell growth in response to 5-30 nM obatoclax (OBT) in PyMT-Bo1 and PyMT-Bo1-abemaR cells at 72 h. (B, C) Cell growth in PyMT-Bo1 and PyMT-Bo1-abemaR cells treated with vehicle, 500nM abemaciclib, 5μM HCQ, 15 nM OBT, or the dual and triple combinations for 72 h. Red dashed line marks the inhibitory effect of the combination of HCQ+abemaciclib in respective cell lines. ***p<0.05 and ****p<0.01 compared to vehicle (Veh); ##p<0.01 compared to HCQ+abemaciclib by one-way ANOVA.

**Figure S7.**
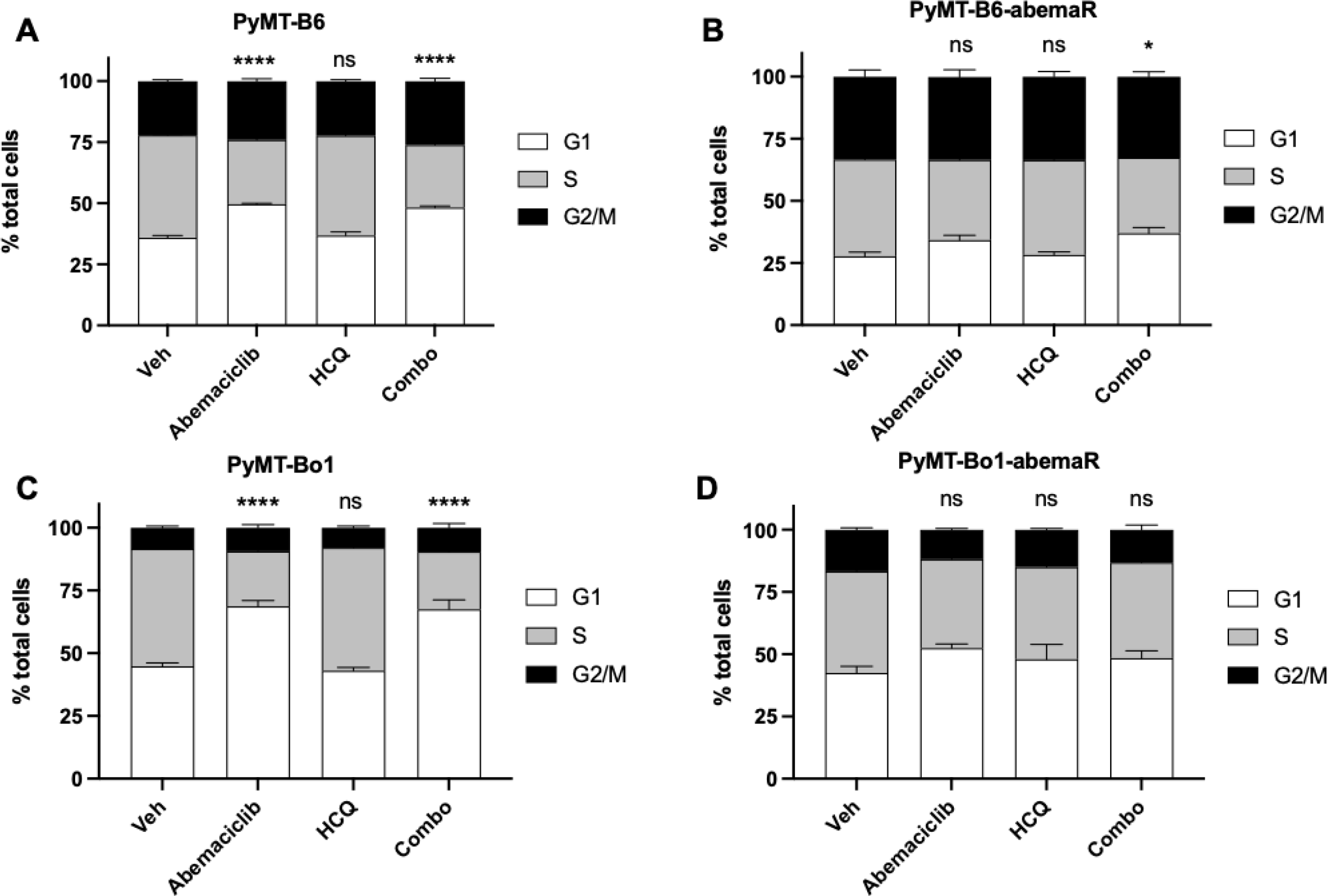
Combination of HCQ and abemaciclib does not augment G1 cell cycle arrest in abemaciclib resistant cells. Cell cycle analysis of (A) PyMT-B6, (B) PyMT-B6-abemaR, (C) PyMT-Bo1, and (D) PyMT-Bo1-abemaR cells after treatment with vehicle, 500nM abemaciclib, 5μM HCQ, or the combination for 72h. *p<0.05; ****p<0.0001; ns, not significant.

**Figure S8.**
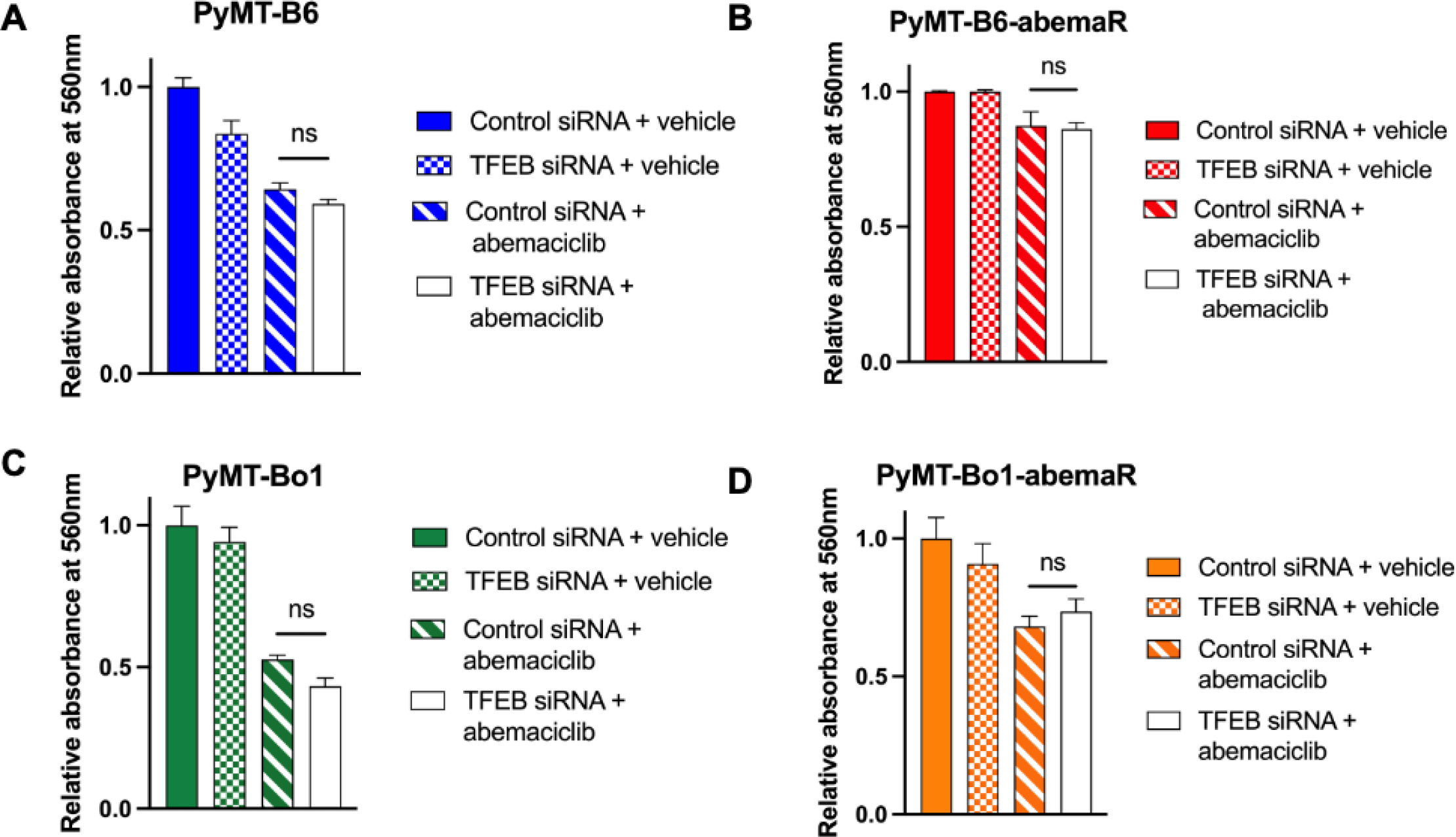
Addition of abemaciclib to TFEB knockdown does not re-sensitize resistant cells to abemaciclib. (A,B) Cell growth determined by crystal violet assays following transfection with control or *TFEB* siRNA and treatment with vehicle alone or 500nM abemaciclib in (A) PyMT-B6, (B) PyMT-B6-abemaR, (C) PyMT-Bo1 and (D) PyMT-Bo1-abemaR cells. ns, not significant by one-way ANOVA.

**Table S1:** Significantly changed lysosomal proteins in abemaciclib resistant PyMT-B6-abemaR cells compared with sensitive PyMT-B6 cells.

| **Protein** | **Average Log2 Ratio**<br>**(resistant/sensitive)** | **Q Value** |
| --- | --- | --- |
| CD68 | 5.69163 | 0.003475 |
| CPQ | 3.525815 | 0.00532 |
| DPP7 | 2.391371 | 0.032419 |
| DNASE2 | 2.312385 | 0.043787 |
| ARSB | 2.098947 | 0.002511 |
| FUCA1 | 2.007189 | 0.02041 |
| BACE1 | 1.952386 | 0.012021 |
| SGSH | 1.847237 | 0.025798 |
| GALC | 1.744517 | 0.008219 |
| PDGFRB | 1.61247 | 0.009095 |
| SMPD1 | 1.595341 | 0.008228 |
| CTSZ | 1.546014 | 0.000505 |
| GALNS | 1.478873 | 0.016925 |
| CTSA | 1.430706 | 0.018544 |
| GM2A | 1.364175 | 0.017459 |
| PLBD2 | 1.24979 | 0.012675 |
| HEXB | 1.221419 | 0.027923 |
| CHID1 | 1.204597 | 0.00833 |
| AGA | 1.088881 | 0.021961 |
| MANBA | 1.088299 | 0.030497 |
| HGSNAT | 1.084099 | 0.045413 |
| LMBRD1 | 1.07601 | 0.039157 |
| ADA | 1.057536 | 0.007594 |
| NPC2 | 1.050028 | 0.023793 |
| CTBS | 1.024642 | 0.010376 |
| HEXA | 1.013703 | 0.031301 |
| PPT1 | 0.997935 | 0.014765 |
| GLB1 | 0.995372 | 0.025238 |
| NAGLU | 0.9861 | 0.033439 |
| ABCA2 | 0.92055 | 0.004578 |
| CCZ1 | 0.89681 | 3.60E-05 |
| PCYOX1 | 0.864156 | 0.005451 |
| TPP1 | 0.821853 | 0.041004 |
| ASAH1 | 0.813657 | 0.009352 |
| TMEM9 | 0.796506 | 0.001773 |
| GAA | 0.742536 | 0.043787 |
| GNS | 0.728895 | 0.03431 |
| SLC48A1 | 0.714791 | 0.026456 |
| VPS13C | 0.70775 | 2.22E-05 |
| CTSD | 0.701908 | 0.031979 |
| BRI3 | 0.699094 | 0.014841 |
| ARL8A | 0.62684 | 0.042056 |
| GGH | 0.616062 | 0.017045 |
| TMEM165 | 0.607021 | 0.022283 |
| RB1CC1 | 0.595083 | 0.000728 |
| ARSA | 0.033306 | 0.033306 |
| UBXN6 | -0.618949 | 6.12E-05 |
| PIP4K2A | -0.658105 | 0.026123 |
| IFITM3 | -0.970928 | 0.018943 |
| CRYAB | -2.045718 | 0.000705 |
| CTSH | -2.112447 | 3.07E-06 |

**Table S2:**
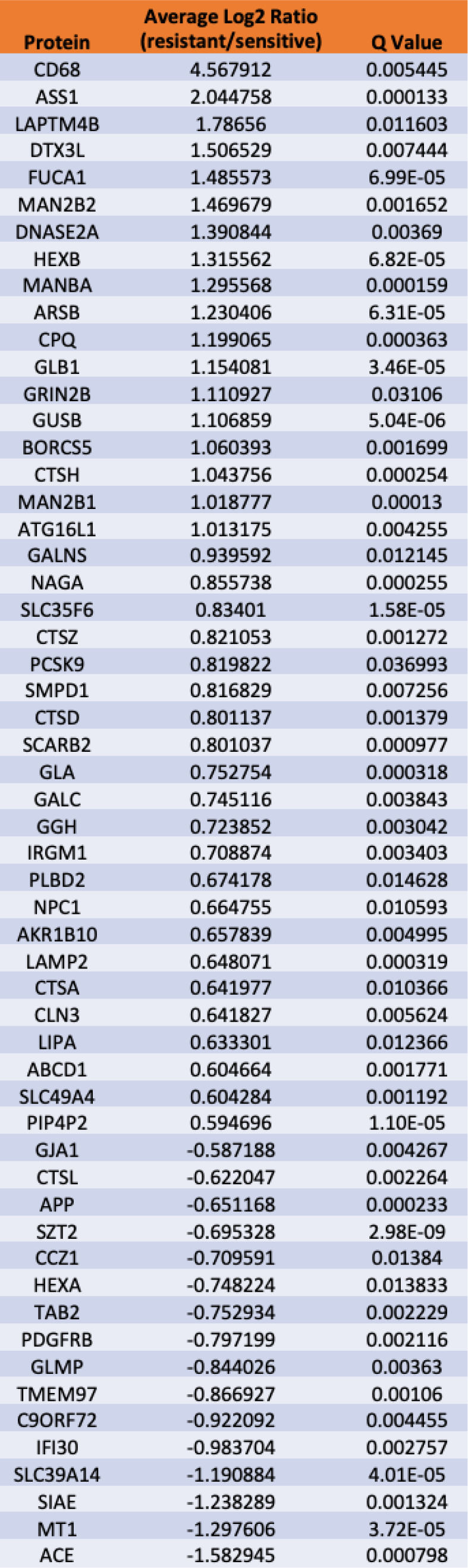
Significantly changed lysosomal proteins in abemaciclib resistant PyMT-Bo1-abemaR cells compared with sensitive PyMT-Bo1 cells.

**Table S3:** Significantly changed lysosomal proteins in MCF7 treated with 1μM abemaciclib or vehicle alone.

| Protein | Average Log2 Ratio<br>(abemaciclib/veh) | Q Value |
| --- | --- | --- |
| LDLR | 1.508403 | 0.001407 |
| SGSH | 1.446615 | 4.40E-05 |
| DPP7 | 1.040809 | 3.33E-10 |
| CTSD | 0.985461 | 6.10E-07 |
| PLBD2 | 0.967666 | 1.44E-09 |
| MAN2B1 | 0.91534 | 0.006602 |
| FYCO1 | 0.890453 | 0.001044 |
| CD63 | 0.886728 | 2.59E-09 |
| BLOC1S1 | 0.820279228 | 0.00453795 |
| CTSL | 0.770206496 | 2.85E-05 |
| PSAP | 0.752527 | 0.000255 |
| LRBA | 0.718388 | 3.27E-07 |
| GAA | 0.712212 | 3.11E-09 |
| SERPINB1 | 0.712114 | 5.62E-07 |
| GNS | 0.698949 | 8.34E-05 |
| PIP4K2A | 0.696472 | 0.020443 |
| LAMP1 | 0.671353 | 2.66E-05 |
| ELAPOR1 | 0.617476 | 5.95E-07 |
| CTSA | 0.617409 | 2.78E-07 |
| SLC7A5 | -0.644423599 | 7.08E-07 |
| SLC29A3 | -0.742482 | 0.036628 |
| SLC35F6 | -1.353814 | 6.20E-05 |

